# The regulator-executor-phenotype architecture shaped by natural selection

**DOI:** 10.1101/026443

**Authors:** Han Chen, Chung-I Wu, Xionglei He

## Abstract

The genotype-phenotype relationships are a central focus of modern genetics. While deletion analyses have uncovered many regulatory genes of specific traits, it remains largely unknown how these regulators execute their commands through downstream genes, or executors. Here, we wish to know the number of executors for each trait, their relationships with the regulators and the role natural selection may play in shaping the regulator-executor-phenotype architecture. By analyzing ∼500 morphological traits of the yeast *Saccharomyces cerevisiae* we found that a trait is often controlled directly by a large number of executors, the expressions of which are affected by regulators. By recruiting a set of “coordinating” regulators, natural selection helps organize the large number of executors into a small number of co-expression modules. This way, the individual executors can be readily recognized by observational approaches that examine the statistical association between gene activity and trait. When the trait is subject to little or no selection, however, the executors are controlled only by “non-coordinating” regulators that evolve passively and do not build the executors’ co-expression. As a result, none of the executors remain a statistically tractable relationship with the trait. Thus, natural selection by governing some traits strongly (such as fertility) and others weakly (such as aging-related phenotypes) profoundly influences the genotype-phenotype relationships as well as their tractability.

## Introduction

Characterization of genotype-phenotype relationships relies on two basic strategies: observational approaches that consider the statistical association between gene activity and a trait and perturbational approaches that consider the mutational effect of a gene on a trait. It is unclear, however, under what circumstances they can be most successful and when they are doomed to fail. Technical advances in recent years enable genomic profiling of various types of gene activity (e.g., mRNA level, protein abundance, protein phosphorylation, protein location, protein-protein interactions, protein-DNA/RNA interactions)(1), greatly facilitating the observational approaches to inferring gene-trait associations. As for the perturbational (or genetic) approaches, it has been well demonstrated that the genome-wide association study as a forward genetic approach is highly powerful in associating DNA mutations/variants to a trait(2); meanwhile, the emerging genome-wide reverse genetic screenings based on homologous recombination(3), RNAi(4) or CRISPR-Cas9(5) are designed to reveal the whole set of genes whose loss-of-function mutations can change a trait. It is thus increasingly clear that data acquisition is no longer a major hurdle to understanding the genotype-phenotype relationships. However, three key challenges remain. First, the performance of observational approaches is heavily compromised by between-gene epistasis that appears to be pervasive(6). Second, because all genes are connected with each other in a cell to influence traits, mutating a random gene could, in principle, propagate through the cellular system to affect every trait to some extent(7, 8). A conceivable scenario is to, say, mutate the gene encoding beta-actin, which likely alters all cellular processes as well as all traits. Because no functional insight can be gained from claims of a gene affecting all traits or a trait affected by all genes, it is unclear when and how perturbational approaches can provide trait-specific functional information. Third, there is no framework to integrate the genes revealed by observational approaches with the genes revealed by perturbational approaches(9).

There issues could be clarified if a general gene architecture underlying a trait was available. A previous study analyzed the microscopic images of triple-stained cells of the yeast Saccharomyces cerevisiae and characterized quantitatively 501 morphological traits for each of 4,718 yeast mutants each lacking a different non-essential gene(10). And, ∼1,500 of the yeast single-gene deletion mutants have microarray-based expression profiles generated recently(11). We can combine the two large datasets to study the relationships of gene deletion, gene expression and trait. In addition, because some of the 501 traits are tightly coupled with fitness while some are decoupled from fitness, there is a unique opportunity to assess the role of natural selection on shaping the genotype-phenotype relationships. We show that the gene architecture underlying a complex trait is often hierarchical with two layers: the lower layer comprises a large number of “executor” genes that are directly responsible for the trait; the upper layer comprises “regulator” genes that control the expressions of the executors. Significant genetic effects usually results from deleting a regulator instead of an executor because more executors are affected in the former. Executors are recognized by observational approaches but they are, due to the constraint imposed by statistics, recognizable only when they form a small number of co-expression modules. Regulators are recognized by perturbational approaches, with two distinct types: “coordinating” regulators that are responsible for the executors’ co-expression and “non-coordinating” regulators that do not contribute to executors’ co-expression. Importantly, the coordinating regulators must be recruited and/or maintained by natural selection to govern fitness-coupled traits, so neither observational nor perturbational approaches succeeded in revealing the executors of a trait subject to little selection. These results illustrate vividly how nature selection acts to build the order of a living system, which ensures the tractability of the system. Without the protection of selection the genes are inevitably disorganized, rendering enormous confusions in the research practice that are discussed in the end of the article.

## Results

### Observational approaches target executors while perturbational approaches target regulators

We used mRNA level as the representative gene activity to test observational approaches in revealing the genes associated with a trait. For each of the 501 yeast traits we identified the genes whose expression is significantly correlated to the focal trait in a robust fashion (Methods). These genes are termed as <u>*o*</u>bservationally <u>*i*</u>nformative <u>*g*</u>enes (OIGs). The proportion of trait variance explained by an OIG is 3.4±2.1% (mean±s.d.). The number of OIGs found in a trait varies substantially across the traits, ranging from zero to ∼1,000. A total of 2,541 non-redundant yeast genes (>40% of all genes) are identified as executors in at least one trait, with the mean and median number of traits an executor affects being 27 and 11, respectively. Some of the OIGs may causally affect their traits (causal OIGs) while the others may be reactive to their traits (reactive OIGs). To estimate the proportion of causal OIGs, we examined the same expression-trait correlations of 118 OIGs that are also detected in ∼60 F1 segregants of a hybrid of two *S. cerevisiae* strains(12). Following a previously proposed method(13, 14), we analyzed the segregation pattern of trait, gene expressions and quantitative trait loci (QTL) in the F1 segregants and estimated that ∼40% of the OIGs are causal (Methods) (Fig. 1A).

**Fig. 1.**
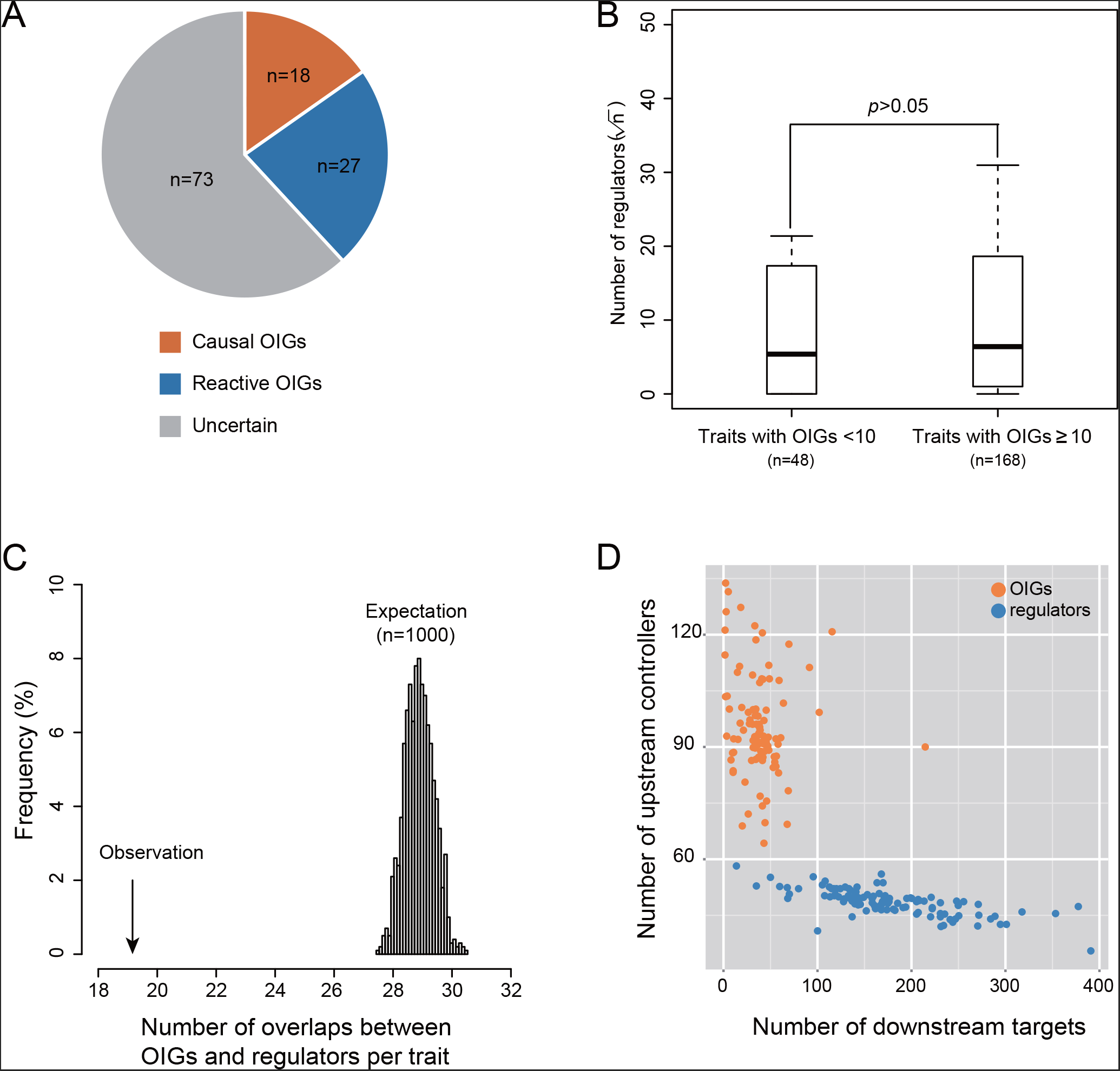
Comparisons between the OIGs revealed by observational approaches and the regulators revealed by perturbational approaches. **(A)** Estimation of the proportion of causal OIGs. **(B)** Comparable numbers of regulators in traits with distinct numbers of OIGs. Box-plots are presented, with the y-axis showing the square root of the regulator number. Mann-Whitney U test is used to compute the *p* value. **(C)** The observed overlaps between OIGs and regulators of the same trait are fewer than the expected, which is estimated by reshuffling the OIGs and regulators of the same trait. The average number of overlaps per trait is presented, and the distribution of expectation is derived from 1,000 independent reshuffles. **(D)** The number of downstream targets (x-axis) and upstream controllers (y-axis) per executor (or regulator). Each dot represents a trait, and the average of all OIGs (or regulators) of the trait is presented. Gene A is called the downstream target of gene B and B the upstream controller of A, if gene A shows a significant expression change (*p* < 0.0001) in the mutant lacking gene B.

The genes whose deletions result in statistically significant effects on a trait are termed as regulators of the trait. We considered the regulators defined by a previous study for 216 out of the 501 yeast traits under a stringent statistical cutoff(15) (Methods). Presumably, a trait with no or just a few OIGs might be a simple trait with a limited number of regulators. This is, however, not true because there are on average 129±22.9 regulators for traits with <10 OIGs and 167±15.6 for the rest traits with ≥ 10 OIGs (*p* > 0.05, Mann-Whitney U test; Fig. 1B). Analysis of the 109 traits each with ≥ 10 OIGs and ≥ 10 regulators shows that OIGs tend not to be the regulators of the same traits (*p* < 0.001, Permutation test; Fig. 1C). Gene Ontology (GO) analysis of the 200 most pleiotropic regulators shows that up to 63.5% of them have the GO term of “Biological Regulator” (*q* < 1.2x10^−16^), while no strong GO enrichment is found for OIGs. These results suggest that the observational approach and perturbational approach are complementary, likely targeting different parts of the gene architecture of a trait. Analysis of expression data shows that, compared to deleting an OIG, deleting a regulator tends to affect a far larger number of downstream genes, while regulators have far fewer upstream controlling genes than OIGs (Fig. 1D). A reasonable explanation to the pattern is that a typical complex trait is controlled directly by a large number of OIGs such that the effect of deleting a single OIG is often too small to be detected; large genetic effects that can be easily detected often result from deleting a regulator, which changes many OIGs simultaneously. This is analogous to a production line run by workers who are supervised by managers, where major productivity slow-down is often due to removing a manager instead of removing a worker although the workers are directly involved in production. Thus, the gene architecture underlying a typical complex trait appears to be hierarchical with two layers: the lower layer comprises a large number of executors that are directly responsible for the trait, and the upper layer comprises regulators that control the activity (expression) of the executors. Executors are supposed to be revealed by the observational approach as OIGs. For convenience hereafter OIGs are termed as observed executors although only ∼40% of them are estimated to be true executors (i.e., causal OIGs). Regulators are revealed by the perturbational approach; because traits and gene expressions are measured in the same gene-deletion mutants, regulators of a trait are also regulators of the executors of the trait.

### Executors are recognizable only when they are co-expressed as a result of natural selection

It is intriguing that the observational approach performed so differently, revealing up to ∼1,000 executors in some traits but zero in some others. On the one hand, despite the contamination of false positives it is still surprising to observe in a single trait as many as ∼1,000 executors each explaining ∼3.4% of the trait variance, because the trait variance explained by each causal factor will be negligible if the number of independent causal factors is large. A reasonable explanation is that the large number of executors are organized into a small number of co-expression modules, which is supported by the data (Fig. S2). On the other hand, many traits have virtually no observed executors although they have as many regulators as the others (Fig. 1B). One possibility is that they indeed have a very limited number of executors. If this is true, such executors should each explain a large proportion of the trait variance and are readily observed. We examined the 54 traits each with 1-3 observed executors and found that the trait variance explained by the observed executors is invariably small (Fig. S1). The remaining explanation is that there are actually a large number of executors underlying each of the traits. Observational approaches failed to reveal them because they are not co-expressed such that the trait variance explained by each individual executor is too small to be detected. This idea, however, cannot be directly tested because the executors are mostly unknown. Because executors of a trait in general should have a stronger expression-trait correlation than non-executors, in each trait we selected the top 100 genes with the strongest expression-trait correlation and calculated the pairwise co-expression levels of the 100 genes. The number of observed executors of a trait appears to be well explained by the average co-expression level of the selected top 100 genes of the trait (Spearman’s ρ = 0.76, n = 501, *p* < 10^−16^; Fig. S2).

The hypothesis that the performance of observational approaches is determined by the executors’ co-expression requires a more critical test. We reasoned that in a biological system such co-expression must be built or maintained by natural selection, so it can be strong only in the traits tightly coupled with fitness. If the co-expression hypothesis is correct, the observational approach should succeed the most in the fitness-tightly-coupled traits and fail in the traits decoupled from fitness. We used cell growth rate as a proxy of fitness, which is reasonable for the single-celled yeast(3), and calculated the trait relatedness to fitness for each of the 501 trait (Methods). The larger absolute value of a trait relatedness to fitness, the stronger coupling of the trait with fitness. Remarkably, the number of observed executors of a trait is largely explained by the trait relatedness to fitness (Spearman’s ρ = 0.89, n = 501, *p* < 10^−16^). There are typically several hundred observed executors in a fitness-tightly-coupled trait but no observed executors at all in the traits with no significant correlation to fitness (Fig. 2). This pattern remains when 57 largely unrelated exemplar traits are considered (Methods) (Fig. S3A). Because the common expression responses to fitness reduction may confound the analysis, we excluded from the analysis all executors observed in more than one exemplar trait and obtained a similar pattern (Fig. S3B). Also, the pattern cannot be explained by the noise in trait measuring (Fig. S4) or by a smaller trait variation in fitness-less-coupled traits (Fig. S5). The fact of no observed executors in most fitness-uncoupled traits and the two orders of magnitude difference between the fitness-coupled and fitness-uncoupled traits lend strong support to the co-expression hypothesis, even after considering the false positives in the observed executors. In summary, for a trait with a large number of executors, observation approaches can succeed only when the executors are organized by natural selection into a small number of independent working modules; the lack of selective constraint results in poor co-expression of the executors such that none of the individual executors alone explain a statistically tractable proportion of the trait variance. Note that for simplicity throughout the manuscript fitness-coupled (-uncoupled) traits refer to those whose trait value is tightly (loosely) coupled with cell growth rate. We are fully aware that all traits are fitness-coupled to some extent.

**Fig. 2.**
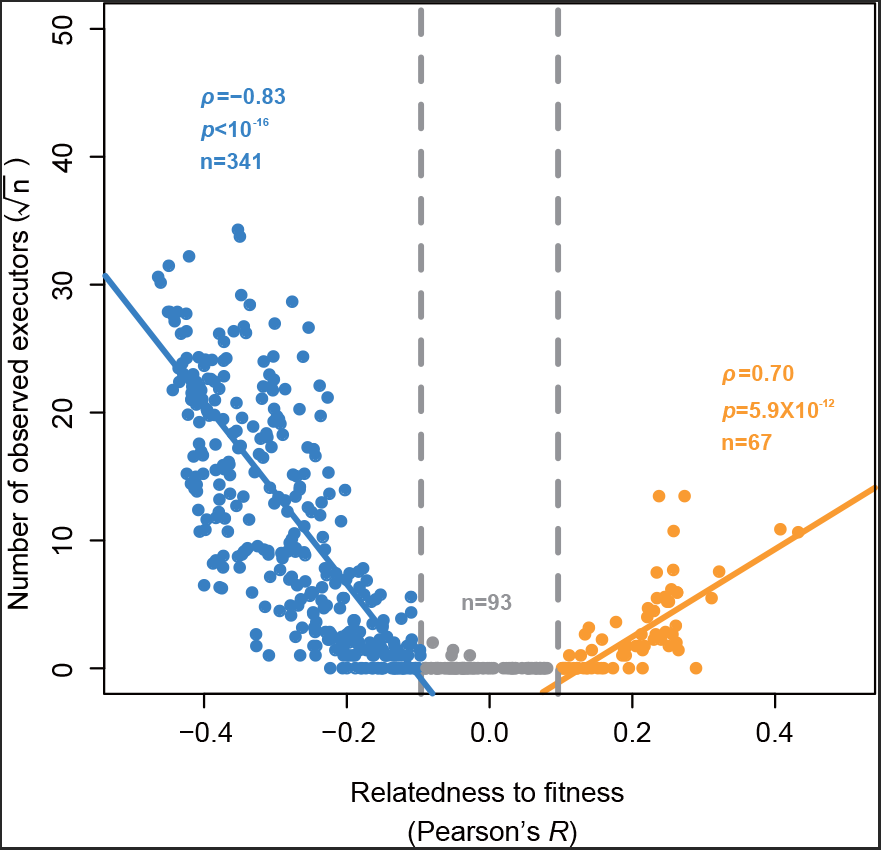
The performance of observational approaches in a trait is determined by the trait relatedness to fitness. The y-axis shows the square root of the number of observed executors, and the x-axis is the trait relatedness to fitness measured by the Pearson’s *R* between the trait values and the cell growth rates of the yeast mutants. Larger absolute *R* means stronger fitness coupling, and -0.096 < *R* < 0.096 corresponds to the statistically insignificant range after controlling for multiple testing. Each dot represents a trait, and ρ shows the Spearman’s correlation coefficient.

### The co-expression of executors is realized by coordinating regulators

Mechanistically, the co-expression of executors is realized by regulators. But, not all regulators contribute to the co-expression, as evidenced by the on average >100 regulators present in most fitness-uncoupled yeast traits in which such co-expression does not exist (Fig. S6). We reasoned that some regulators of a given trait may help build the executors’ co-expression (coordinating regulators) while the others may affect the executors non-concordantly and destroy such executors’ co-expression (non-coordinating regulators). Coordinating regulators are supposed to be recruited and/or maintained by natural selection; and, to achieve a strong co-expression, they should regulate as many as the executors; most importantly, their across-executor regulation profiles should be congruent with each other (Fig. 3A). Non-coordinating regulators, however, do not have to be congruent with the coordinating regulators or with each other in their regulation profile (Fig. 3A). Also, their effects on the executors may evolve passively(16, 17), for some other purposes or just by neutral drift, which is highly likely considering the dense transcription factor bindings sites on a gene(18) and the widespread gene pleiotropy(19). Because deletion of a coordinating regulator launches a coordinated attack on many of the executors, the effect should be in general much larger than that of deleting a non-coordinating regulator. This prediction is helpful for identifying coordinating regulators.

**Fig. 3.**
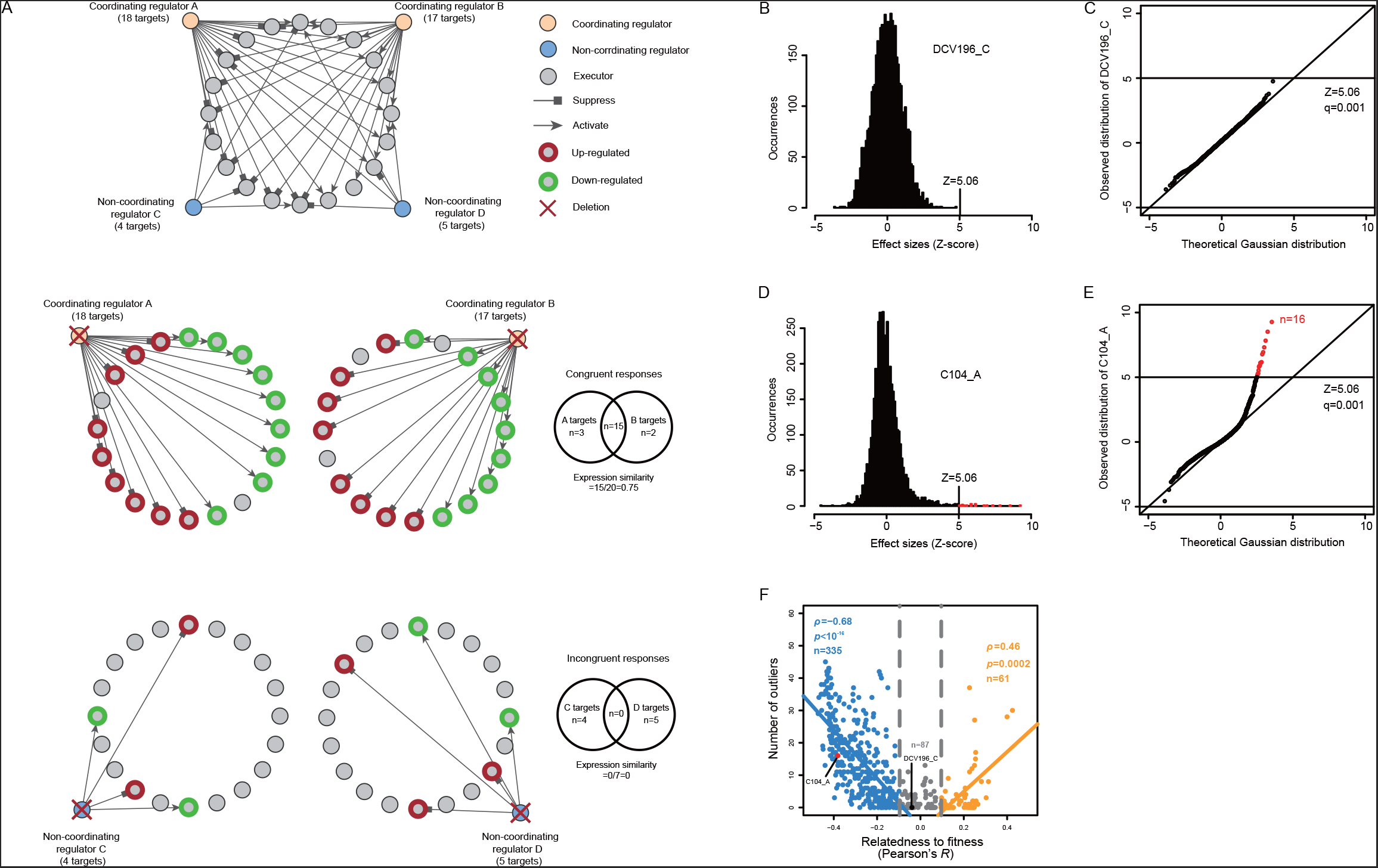
Characterization of coordinating and non-coordinating regulators. **(A)** Proposed features of coordinating and non-coordinating regulators. The across-executor regulation profile is congruent for different coordinating regulators of the same trait, which is, however, not the case for non-coordinating regulators. **(B, C)** The frequency distribution of the normalized trait values (Z-score effect sizes) and the Q-Q plot that compares this distribution with the Gaussian approximation for trait DCV196_C. **(D, E)** Same as the panels B and C, except for trait C104_A. **(F)** The number of outliers found in a trait is well explained by the trait relatedness to fitness. Larger absolute *R* means stronger fitness coupling, and -0.096 < *R* < 0.096 corresponds to the statistically insignificant range after controlling for multiple testing. Each dot represents a trait, and ρ shows the Spearman’s correlation coefficient.

For each trait we modeled the frequency distribution of effect size upon deleting each of the 4,718 yeast genes using the Gaussian function that is commonly used to model a quantitative trait(20) (Methods). We used quantile-quantile plot to compare the Gaussian approximation to the true distribution and found that the two distributions often fit each other reasonably well (Fig. 3B, C). In some traits, however, there are disproportionately large effects that are far beyond the Gaussian approximation (Fig. 3D, E). In each trait we defined outlier effects as those with the absolute effect size Z-score > 5.06, which corresponds to *p* = 2.12 x 10^−7^ in the standard Gaussian distribution or *q* = 0.001 after the Bonferroni correction for multiple testing (*q = p* x 4,718). The number of outlier regulators found in a trait ranges from zero to ∼50. If the outliers represent coordinating regulators, they should be present primarily in fitness-coupled traits because selection is required to recruit/maintain them. Indeed, the number of outliers found in a trait is well explained by the trait relatedness to fitness (Spearman’s ρ = 0.85, n = 483, *p* < 10^−16^) (Fig. 3F). This pattern remains when the Z-score cutoff of defining outliers is changed to 4.56 (*q* < 0.005) or to 4.06 (*q* < 0.01) (Fig. S7), or when only the exemplar traits are analyzed (Fig. S8). Because this correlation could be explained by the possibility that the presence of outliers causes strong fitness coupling, we recalculated for each trait the relatedness to fitness after excluding the outlier genes (Methods). The recalculated values are highly correlated to the original ones (Pearson’ *R* = 0.96, n = 501, *p* < 10^−16^; Fig. S9). Thus, the correlation is better explained by the opposite scenario that the strong fitness coupling of a trait helps recruit and/or maintain the outliers found in the trait. This analysis suggests that the outliers might be the coordinating regulators we are searching for.

If this is true, we expect plenty of common expression responses upon deleting different outliers of the same trait because coordinating regulators of a trait are characterized by their congruent regulation profiles with each other (Fig. 3A). Such common expression responses are not expected when non-coordinating regulators are deleted because in that context the executors are altered non-concordantly. To make a fair comparison in each trait we examined the top 20 regulators with the largest deletion effect as well as available expression information (Methods). The proportion of outliers among the selected 20 regulators of a trait ranges from zero to 90%, depending on the trait relatedness to fitness (Fig. 4A). For each trait we identified the genes that are commonly up- or down-regulated among the mutants of the 20 regulators, under a statistical cutoff where the expected number of such genes is slightly smaller than one (0.6). The number of such commonly responsive genes of each trait is highly dependent on the proportion of outliers (Spearman’s ρ = 0.74, n = 129, *p* < 10^−16^; Fig. 4B), while the total number of responsive genes in the 20 mutants is similar across the different traits (Spearman’s ρ = 0.07, n = 129, *p* = 0.43; Fig. S10). Importantly, most of the commonly responsive genes are also the observed executors of the same traits (60%; 8.7-fold enrichment with the 95% confidence interval of 5.6∼12.2-fold; permutation test). Thus, the outliers found in a trait are indeed coordinating regulators of the trait, with congruent profiles in regulating the executors. This being said, we note that both false positives and false negatives must exist: some weaker coordinating regulators may show a non-outlier deletion effect and some outlier effects may arise from the non-coordinating regulators that happen to affect a lot of executors or some large-effect executors.

**Fig. 4.**
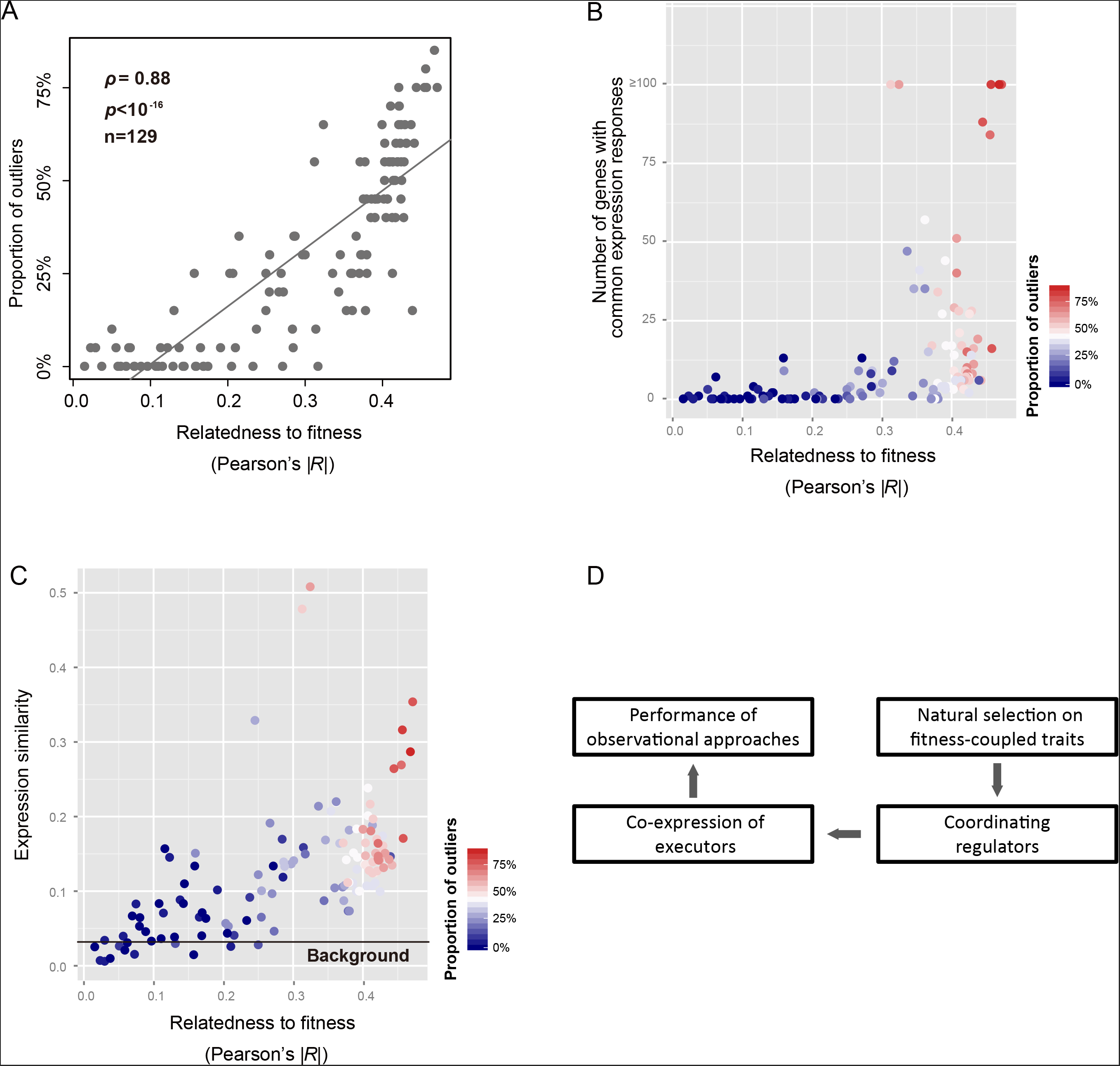
Outliers found in a trait are coordinating regulators with congruent regulation profiles. **(A)** The proportion of outliers among the 20 selected regulators of a trait is a function of the trait relatedness to fitness. Each dot represents a trait. **(B)** The number of commonly responsive genes upon deleting the selected regulators of a trait is positively correlated to the proportion of outliers among the selected regulators of the trait (Spearman’s ρ = 0.74, n = 129, *p* < 10^−16^) or the trait relatedness to fitness (Spearman’s ρ = 0.65, n = 129, *p* < 10^−16^). Each dot represents a trait. **(C)** The expression similarity among all non-outlier mutants in a fitness-uncoupled trait is similar to the background between-mutant expression similarity. Expression similarity for a trait is the average Pearson’s *R* of all 190 pairs of the mutants of the 20 selected regulators. The background shown as a bold line is the average Pearson’s *R* of all pairs of the mutants of the selected regulators across the 129 traits. Each dot represents a trait. **(D)** The logic chain revealed by this study.

An equally important observation is the nearly zero commonly responsive genes (mean=1.46± 2.23, median=0, n=28) found in the traits with 100% non-outlier regulators that are primarily non-coordinating. To see whether this signal is robust, we considered the genome-wide expression similarity among the 20 selected mutants of each trait (Methods). The resulting expression similarity of a fitness-uncoupled trait with all non-outliers is no stronger than the background, which is the average expression similarity among all selected mutants of the 129 traits examined here (Fig. 4C). This result indicates that deleting non-coordinating regulators provide little or no trait-specific functional information even if the resulting genetic effects are the strongest on the trait, since the selected regulators are all the strongest ones in their own trait. It also explains from another angle why we failed to observe any executors in fitness-uncoupled traits.

Therefore, a complete logic chain summarizing all the analyses is available: For each trait the performance of observational approaches is determined by the executors’ co-expression, which is realized by coordinating regulators that are recruited and/or maintained by natural selection, a process being effective only when the trait is fitness-coupled (Fig. 4D).

### Natural selection but not genetic heterogeneity determines the success of modeling a trait

The idea of using observed executors to model a trait has been common(21), but the performance is not stable(22). Although there could be many confounding issues(1), the complexity of genetic architecture seems to be the rate-limiting factor. To test the upper limit of modeling a trait with observed executors, we chose to examine cell growth rate, the yeast fitness-determining trait with arguably the most complex genetic architecture since over one third (∼2,000) of the yeast genes, when deleted, show a growth rate reduction greater than 5% in the rich medium YPD(23). Using the functional data considered above, we identified ∼900 observed executors for the growth rate under a stringent criterion. With the help of protein-protein interactions we obtained six modules composed of ∼300 observed executors (Methods), and found that each module is responsible for a critical biogenesis process (Table S1). We examined 87 mutants each with a growth rate less than 80% of the wild-type, and computed the expression distance (ED) between a specific mutant and the wild-type in each of the six modules (Methods). With only a few exceptions, the 87 mutants formed five distinct clusters, each corresponding to the expression alteration of a unique set of modules (Fig. 5A). Because expression responses to slow growth should be conserved among slow-growth mutants, this pattern indicates that the expression alteration of the modules causes the slow growth of these mutants. In other words, the six modules are true executor modules that causally affect the yeast cell growth. Further analysis reveals new understandings on the function of ribosome in affecting the yeast cell growth (Fig. S11). Moreover, a simple linear function integrating the six executor modules explains up to ∼50% of the growth rate variation among over 400 single-gene deletion mutants (Pearson’s *R* = 0.69, n = 443, *p* < 10^−16^; Fig. 5B). Note that the cell growth rates considered here are measured using Bar-seq(23), which is believed more accurate than the microarray-based method(24) or colony-size-based method(6), both used for quantifying the yeast growth rates. For the same mutants, the Pearson’s *R* is 0.77 between the microarray-based measures and the Bar-seq-based measures (Fig. 5C), and 0.63 between the colony-size-based measures and the Bar-seq-based measures (Fig. 5D). This suggests that the executor-based linear model is comparable to the two conventional experimental approaches in estimating the yeast cell growth. The success in such a complex trait suggests that the heterogeneity of genetic architecture is not a major hurdle to modeling a trait. Instead, the level of fitness coupling, which determines to what extent the executors are observable, appears to be the rate-limiting factor. Thus, one should gauge the evolutionary relevance/importance of a biological phenomenon before any attempts to understand the phenomenon, a notion with particularly important implications to all kinds of artificial phenomena invented in laboratory.

**Fig. 5.**
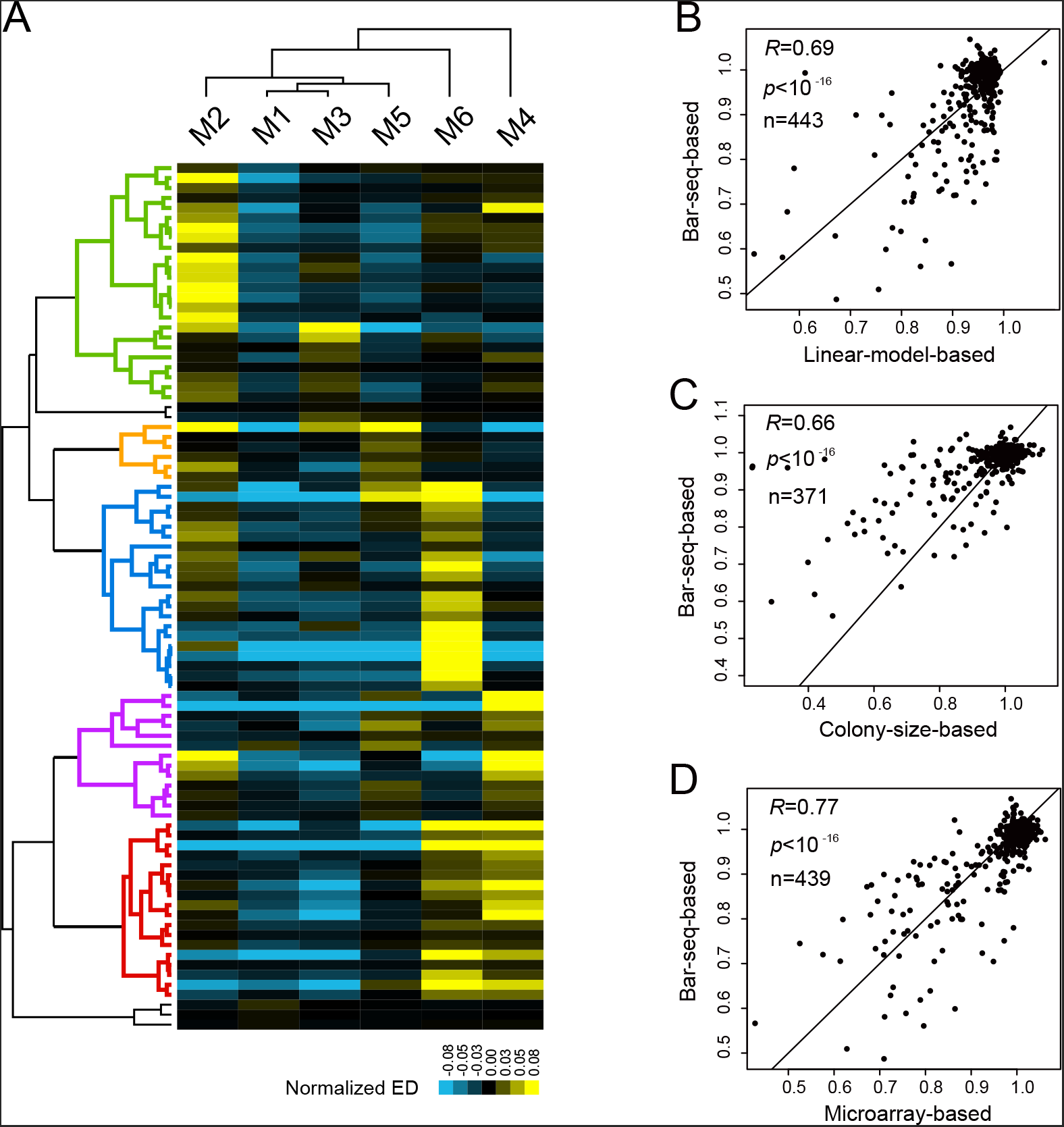
Good performance of the observed executors in modeling a super-complex trait. **(A)** Five types of growth defects defined by the six executor modules. Each row represents a slow-growth mutant, and the expression distance (ED) of a module is normalized by subtracting the average ED of the six modules of the mutant. **(B)** Growth rates of the yeast mutants based on the Bar-seq technique or the linear model written as *G* = -1.740ED_*M1*_ – 0.435ED_*M2*_ – 0.725ED_*M3*_ – 0.071ED_*M4*_ + 0.794ED_*M5*_ -0.058ED_*M6*_ + 1.019, where *G* stands for growth rate and ED_*Mi*_ stands for expression distance of an executor-module away from the wild-type. Each dot represents a single-gene deletion mutant, with the Pearson’s *R* shown. **(C)** Growth rates of the yeast mutants measured by the Bar-seq technique or the colony-size-based method, with 72 mutants excluded due to the lack of the colony-sized-based measures. **(D)** Growth rates of the yeast mutants measured by the Bar-seq technique or the microarray-based method, with four mutants excluded due to the lack of the microarray-based measures.

## Discussion

There are two caveats that warrant discussion. First, because of data availability we used mRNA level as the representative gene activity and studied only morphological traits in the single-celled organism yeast. There are two main conclusions: 1) For a trait controlled by a large number of executors, the executors can hardly be recognized unless they are organized into a small number of causal modules. 2) The coordination of executors requires trans-factors (coordinating regulators) that can be recruited and/or maintained only by selection. The first is based on a statistical principle and the second represents a principle of evolution. Thus, our findings should be of general meaning, although modification/optimization of the details might be required when they are applied to other systems. For example, the coordination may occur across different types of gene activity, or there could be complex regulations within regulators or even within executors. Second, the observed executors are not all true executors. But this concern is alleviated by the strong signals in the comparisons between the fitness-coupled and -uncoupled traits as well as the sheer absence of observed executors in most fitness-uncoupled traits. Because the genetic/phenotypic space represented by the F1 segregants of the BY x RM hybrid is limited, only 118 observed executors were tested for their causal effects on the traits, which gave a rough estimation of the proportion (∼40%) of being true. Sampling more variations in natural populations would give a more accurate estimation, but a refined estimate is unlikely to overturn the argument of a significant proportion of them being true. Notably, this argument is supported by the six biogenesis-related executor modules that were shown to causally affect the yeast cell growth, since these modules are composed of ∼300 genes selected from the ∼900 observed executors of the trait.

The whole points of this study are summarized in Fig. 6. Specifically, a complex trait is often controlled by a large number of executors, the expressions of which are controlled by regulators of two types. The coordinating regulators organize the large number of executors into a small number of co-expression modules such that the individual executors each have a strong relationship with the trait. In sharp contrast, the non-coordinating regulators affect the executors non-concordantly leading to no co-expression of the executors. As a result, the individual executors hardly have a tractable relationship with the trait, predicting the failure of the attempts to uncover them. A central issue here is that the coordinating regulators must be recruited and/or maintained by natural selection while the non-coordinating regulators may evolve under little or no selective constraint. This model highlights evolution as the unifying principle of biology, an assertion that has never been seriously considered by most (cell) biologists who aim to understand mechanistically how genes are organized in a cell to build up phenotypes(25). Practically, it serves as a general framework for addressing the three key challenges raised at the beginning:

**Fig. 6.**
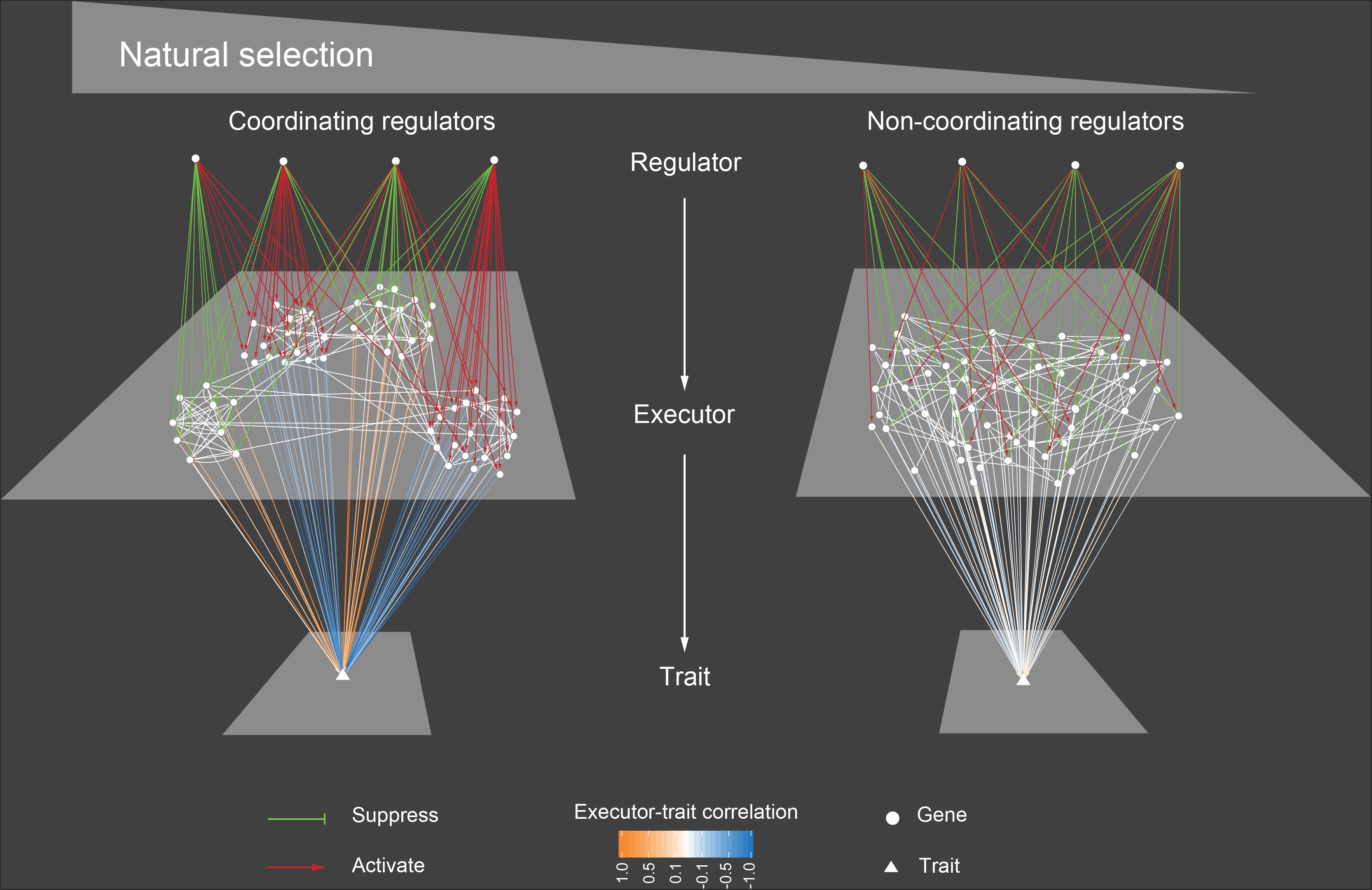
The selection-dependent regulator-executor-phenotype architecture. Natural selection recruits/maintains coordinating regulators to organize the ∼50 executors into four co-expression modules such that each individual executor shows a strong correlation with the trait. When the selection is absent, the ∼50 executors are controlled only by non-coordinating regulators, thus representing ∼50 largely independent causal factors. As a result, none of the executors have a detectable correlation with the trait.

We start with the third challenge, the integration of functional data (by observational approaches) with mutational data (by perturbational approaches). The field has long been puzzled by an observation that the two types of data often do not support each other. For example, the genes with expression responses to a new environment are often not genetically required for dealing with the environment(3, 26). The analogy of the manager-worker combination on a manufacturing line illustrates how the regulator-executor architecture helps reconcile the two types of data.

Another challenge is the functional epistasis that complicates observational approaches. The lack of between-executor coordination predicts that an executor’s effect would always be enhanced or masked by other executors, suggesting pervasive functional epistasis as a result of deregulation(27). Suppose there is a trait controlled by 10 executors each with two functional modes, on or off. The absence of coordination across the 10 executors suggests a total of 2^10^ = 1,024 different functional statuses underlying the trait, so the efforts required for characterizing the 10-gene interactions are simply enormous. This may explain the endless new interactions or pathways discovered in cell biology(28), highlighting the limitation of a purely mechanistic perspective in understanding a cell.

The last challenge is how to interpret genetic effects. A given genetic effect is supposed to (i) suggest a practical means of altering the trait and (ii) provide clues to the functional basis of the trait (i.e., the executors). However, only the former is guaranteed under the current framework of genetics that is defined solely by statistics. Our analysis shows that mutating a non-coordinating regulator of a given trait provides no effective information for revealing the executors because the resulting non-concordant responses of the executors cannot be distinguished from the background expression changes (Fig. 4B and C). Unfortunately, such genetic effects appear to be predominant in number, as up to ∼95% of the statistically significant genetic effects on the yeast traits arise from the deletions of a non-coordinating regulator (Fig. S6). It is likely that many of the confusions in current genetics can be ascribed to the attempts of using non-coordinating regulators to tackle the functional basis of a trait. Recognition of this is helpful for dealing with the pleiotropic effects of a gene(29) and the heterogeneous genetic architecture of a trait (e.g., those called geneticists’ nightmares)(19). Importantly, while natural selection is required to recruit/maintain the coordinating regulators that can help reveal the executors, the relationships between non-coordinating regulators and the trait appear to be of passive origin and subject to no effective selection. We thus argue that, while the century-old statistics-based framework of genetics has been highly successful in Mendelian traits, a selection-based new framework of genetics might be more appropriate for dealing with complex traits.

This study reminds us of a basic biological fact that the order of a living system must be built or maintained by natural selection and the lack of selection renders disorder. The internal order and disorder determine the success of the external research efforts. This notion has special implications to human biology, because selection is inefficient in humans due to the small effective population size(30), and because aging-associated diseases or traits are often of little fitness relevance but of high interest to researchers(31, 32). One may argue that the disordered complexities are exactly the challenges we need to address, but the lack of selective constraint predicts that they might be *ad hoc* phenomena sensitive to genetic and environmental backgrounds(33). A robust discussion of both the necessity and the strategy of studying these issues is needed.

## Materials and Methods

### Data

The yeast *Saccharomyces cerevisiae* singlegene deletion stock was generated by Giaever et al. (2002), with 4,718 mutant strains each lacking a nonessential gene. As for cell growth rates of the 4,718 mutants measured in the rich medium YPD (yeast extract, peptone, and dextrose), the Bar-seq-based data were produced by Qian et al. (2012), the microarray-based data by Steinmetz et al. (2002), and the colony-size-based data by Costanzo et al. (2010). The 501 morphological traits of the mutants were characterized by Ohya et al. (2005) (SCMD). The regulators were defined by Ho and Zhang (2014) under a stringent statistical cutoff for 220 out of the 501 traits; the rest 281 traits contain no information necessary for statistics. We reproduced the analysis and obtained the same set of regulators in 216 out of the 220 traits using the updated data in SCMD. Only the 216 traits are considered in analysis of regulators. The microarray-based expression profiles of 1,484 of the single-gene deletion mutants were generated by Kemmeren et al. (2014); gene A is called the downstream target of gene B and B the upstream controller of A, if gene A shows a significant expression change (*p* < 0.0001 as provided in the original data) in the mutant lacking gene B.

### Identify <u>*o*</u>bservationally <u>*i*</u>nformative <u>*g*</u>enes (OIGs)

For nearly all the morphological traits there is a bell-shape frequency distribution of the trait values among the 4,718 mutants, with the median trait value close to that of the wild-type (Fig. S12). There are 1,328 strains (1325 mutants plus three wild-type strains) each with both the expression profile generated by Kemmeren et al. (2014) and the information of cell growth rate. We randomly divided the 1,328 yeast mutants into two sets, with two thirds (885) for Set #1 and one third (443) for Set #2. There are 6,123 yeast genes on the microarray chip used by Kemmeren et al. (2014). We first generated 500 artificial datasets, each containing 443 strains picked randomly from the 885 Set #1 strains with replacement. We calculated the Pearson’s *R* between expression level and trait value for each of the 6123 genes in each of the 501 traits using each the 500 artificial datasets, respectively. The *R* values were then transformed into *p*-values using t-test, so we obtained 500 *p*-values for each gene in a trait. We defined the correlation robustness (*r*-value) of a gene as the harmonic mean of its *p*-values after dropping both the highest and the lowest 5% of its 500 *p*-values, which was then multiplied by 6,123 for multiple testing correction. Genes with the corrected *r*-values < 0.01 are considered as putative *o*bservationally *i*nformative *g*enes (OIGs). To further reduce false positives, we required that the putative OIGs also show a significant expression-trait correlation in the independent Set #2 mutants. The resulting OIGs are later called observed executors. The trait variance explained by an OIG and the co-expression level between two OIGs are measured in the Set #2 mutants.

Morphological traits are not independent; for instance, the size and the diameter of a cell are correlated. To reduce correlated traits, we used an un-supervised affinity propagation strategy proposed by Frey and Dueck (2007) to cluster the 501 traits based on *r*-values of the 6,123 yeast genes computed in a trait, resulting in 57 clusters each with an exemplar trait.

The frequency distribution of cell growth rates of the yeast mutants is highly biased, with the majority close to the rate of wild-type. We thus computed the expression-growth rate correlation using the univariate Cox’s regression model that emphasizes the difference of two categories, with growth rate as the parameter “time”, strains of growth rate <0.9 weighted as “event = 1”, and all others as “event = 0”. Specifically, we performed the Cox’s regression analysis using the 500 artificial datasets described above and obtained 500 *p*-values for every yeast gene. The corrected *r*-value was computed as described above. A total of 911 genes each with the corrected *r*-value < 0.001 (a more stringent cutoff) were defined as OIGs of the cell growth rate. The 911 OIGs identified in the Set #1 mutants are assembled into protein modules and then tested for their performance in modelling cell growth rates of the 443 Set #2 mutants.

### Define causal and reactive OIGs

Information of the genotype, expression and morphological traits of 62 F1 segregants of a hybrid of two yeast strains (BY4716, a derivative of S288c, and YEF1946, a derivative of RM11-1a) was obtained from Nogami et al. (2007), with three segregants excluded from further analyses because of unmatched IDs. Because no major difference exists between the two parental yeast strains in most of the morphological traits, there are only 118 OIGs whose expression-trait correlations are also observed in the 59 F1 segregants under the false discovery rate of *q* < 0.01 (Bonferroni correction for multiple testing). The causality of the expression-trait associations is resolved using the Network Edge Orienting (NEO) method developed by Aten et al. (2008). Following the manual provided by NEO, we calculated the LEO.NB.CPA score and the LEO.NB.OCA score with all genotype information (SNPs) inputted. For each association the two causality directions (i.e., expression -> trait and trait -> expression) were tested separately. We defined a cause relationship if the LEO.NB.CPA score > 0.8 and the LEO.NB.OCA score > 0.3, which corresponds to a false discovery rate of 0.05. We obtained 18 OIGs each showing expression->trait causal relationship and 27 OIGs showing trait->expression relationship, but failed to assign a reliable causal relationship for the rest 118-18-27=73 associations. Thus, the proportion of causal OIGs is 18/(18+27) = 40% (assuming the same proportion of positives in the 73 uncertain associations).

### Calculate trait relatedness to fitness

Because in this study all cellular traits are measured in YPD, we used the cell growth rate in YPD as the proxy of fitness. Given the bell-shape distribution of a morphological trait where the wild-type trait value is almost always located in the middle, both increase and decrease of a trait value relative to the wild-type could reduce the fitness. Thus, in each trait we divided the 4,718 mutants into two equal halves according to their trait values, and calculated the Pearson’s *R* between trait value and fitness for each half of the mutants separately, resulting in two *R*s for each trait. The half showing the larger *R* (absolute value) is termed as fitness-more-coupled side and the other half as fitness-less-coupled side for the trait. We used the *R* of the fitness-more-coupled side to represent the relatedness to fitness of the trait. To exclude the potential effects of outlier trait values on the estimation of fitness coupling, in each trait we also removed the top 50 trait values from each side and recalculated the trait relatedness to fitness.

### Assess data noise of the 501 traits as well as its potential consequences

To characterize the yeast morphological traits Ohya et al. examined on average 400 individual cells for each mutant. The trait value of a given mutant is the mean trait value of all examined cells. Despite the generally large number of examined cells, in some traits there are only a few tens of informative cells, which may affect the reliability of the measurements. To address this issue, we randomly divided the examined cells of each mutant into two equal halves and computed the morphological traits of each mutant in each half separately. We then compared the trait values obtained from the two halves. The level of consistency between the two halves varies substantially among traits, but this is not dependent on the trait relatedness to fitness.

### Characterize coordinating and non-coordinating regulators

We excluded 18 traits whose effect size distribution is not uni-modal (*p* < 0.05, Hartigan’s Diptest), leaving 483 traits for further analyses. We normalized the raw trait value X_*ij*_ of mutant *j* in trait *i* to Z-score effect size using:

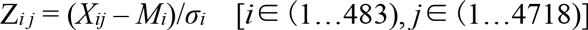

where *M_i_* and *σ_i_* are the mean and standard deviation of the raw trait values in the trait *i* among the 4,718 mutants. The outlier effects are defined as the absolute Z-score > 5.06, which corresponds to *p* < 0.001 x 1/4,718 or *q* < 0.001 according to the standard Gaussian distribution. Accordingly, the regulators with absolute Z-score > 5.06 are outlier regulators (putative coordinating regulators) and the rest are all non-outlier regulators (non-coordinating regulators).

To gain a functional support of the separation of coordinating and non-coordinating regulators, we analyzed the expression profiles of the mutants of these regulators. Out of the 216 traits with regulator information we excluded 20 traits each with outlier regulators in both the fitness-more-coupled and fitness-less-coupled sides of the trait for analysis convenience (there is invariably one outlier regulator in the fitness-less-coupled side for each of the 20 traits). There are 1,325 mutant expression profiles available, so mutants of ∼27% (1325/4718) of regulators can be compared with regard to their expression responses to the deletions of the regulators. For each trait we selected its top 20 regulators with both the largest effect sizes and available expression profiles in the fitness-more-coupled side. We excluded 67 traits each with <20 such regulators. Thus, a total of 216-20-67=129 traits remained for further analyses. For each trait the genes with overall expression up- or down-regulation in mutants of the 20 selected regulators relative to the rest 1,305 mutants were identified as commonly responsive genes in the trait, under the statistical cutoff of *p* < 0.0001 (t-test). There are a total of 1,060 non-redundant yeast genes identified as the commonly responsive genes in at least one of the 129 traits, with the mean and median number of traits a gene involved being 2.97 and 2, respectively. This suggests plenty of trait-specific functional information provided with such common expression responses. Expression similarity between mutants is the average Pearson’s *R* of all pairs of expression profiles of mutants of the 20 selected regulators of a trait. We compared mutants of all selected regulators across the 129 traits to estimate the background between-mutant expression similarity.

### Define executor modules using protein-protein interactions

Yeast protein-protein interactions (PPIs) were downloaded from BioGrid (Stark et al., 2006). We considered only the 911 executors associated with cell growth rate. Protein modules are separated using an order statistics local optimization method (OSLOM) proposed by Lancichinetti et al. (2011) with default settings. We obtained seven modules. To annotate the modules we carried out Gene Ontology (GO) enrichment analysis using BinGO (Maere et al., 2005) and Cytoscape (Shannon et al., 2003). Six modules appear to be highly enriched with functionally similar proteins under a false discovery rate of 0.001 (Table S1).

The expression distance (ED) of an executor module between a mutant and the wild-type is defined as the normalized Euclidian distance:

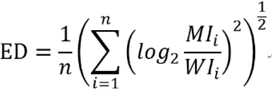

where *MI*_i_ is the expression level of gene *i* in the mutant, *WI*_i_ is the expression level of gene *i* in the wild-type, and *n* is the number of genes in the module.

## Acknowledgments

We are grateful to Drs W. Qian, J. Zhang, X. Liu, P. Shi, R. Zhang, Y. Zhang, W. Zhai, Y. Shen, Y. Huang, T. Tang, B. Liao, S. Xu and M. Bakewell for discussion or comments. X.H. and H.C. designed the study; H.C. and X.H. analyzed data; X.H., H.C. and C. W. wrote the paper.

## Supporting Online Material of “The regulator-executor-phenotype architecture shaped by natural selection”

Han Chen, Chung-I Wu, and Xionglei He*

The State Key Laboratory of Bio-control, College of Ecology and Evolution, School of Life Sciences, Sun Yat-sen University, Guangzhou 510275, China

The SOM contains:

Legends of supplementary figures 1-12

Table S1

Figures S1-12

**Fig. S1.**
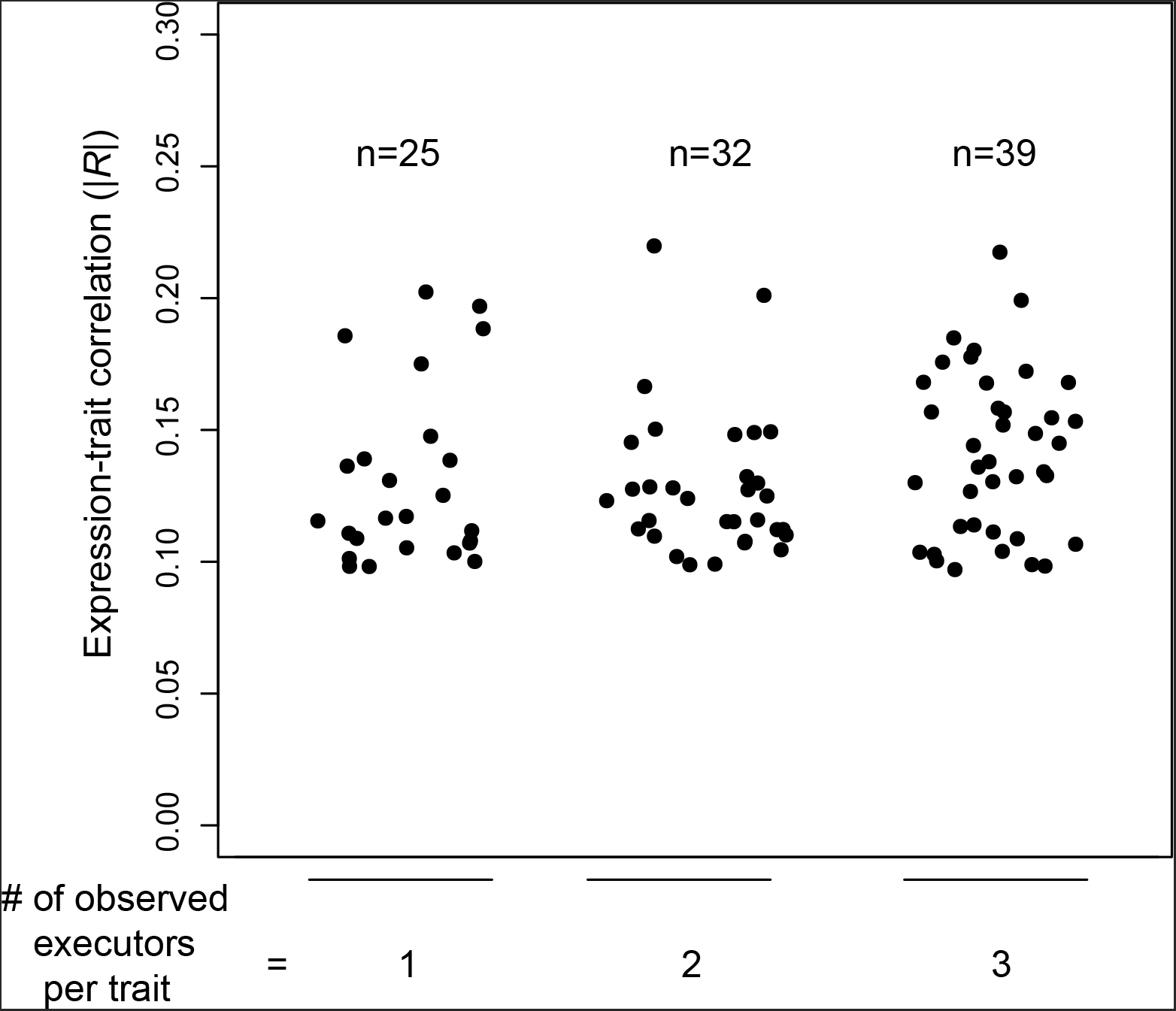
The variance explained by individual observed executors is minimal even in traits with only one, two, or three observed executors (x-axis). The y-axis shows the absolute Pearson’s *R* between gene expression and trait value across the 443 Set #2 mutants. Each dot represents an observed executor.

**Fig. S2.**
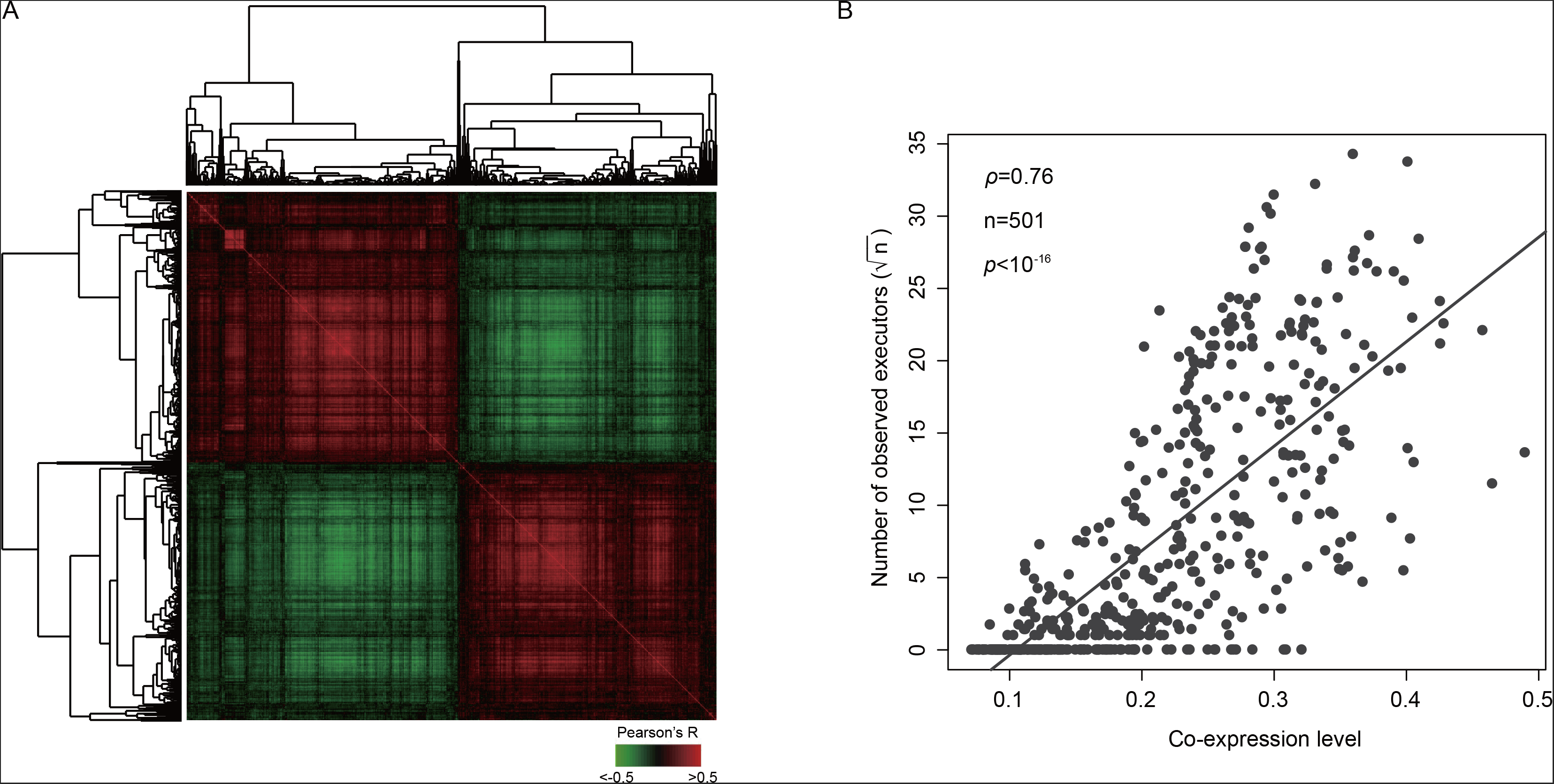
The co-expression level of executors determines their tractability. **(A)** The strong co-expression among the 1,175 observed executors in the trait DCV126-A1B that has the largest number of observed executors. For each gene pair the Pearson’ *R* across the 443 Set #2 mutants is computed. **(B)** The number of observed executors of a trait is highly correlated to the co-expression among the selected top 100 genes of the trait. For each gene pair the absolute Pearson’s *R* across the 443 Set #2 mutants is computed, and the average *R* of all gene pairs is the co-expression level of a trait.

**Fig. S3.**
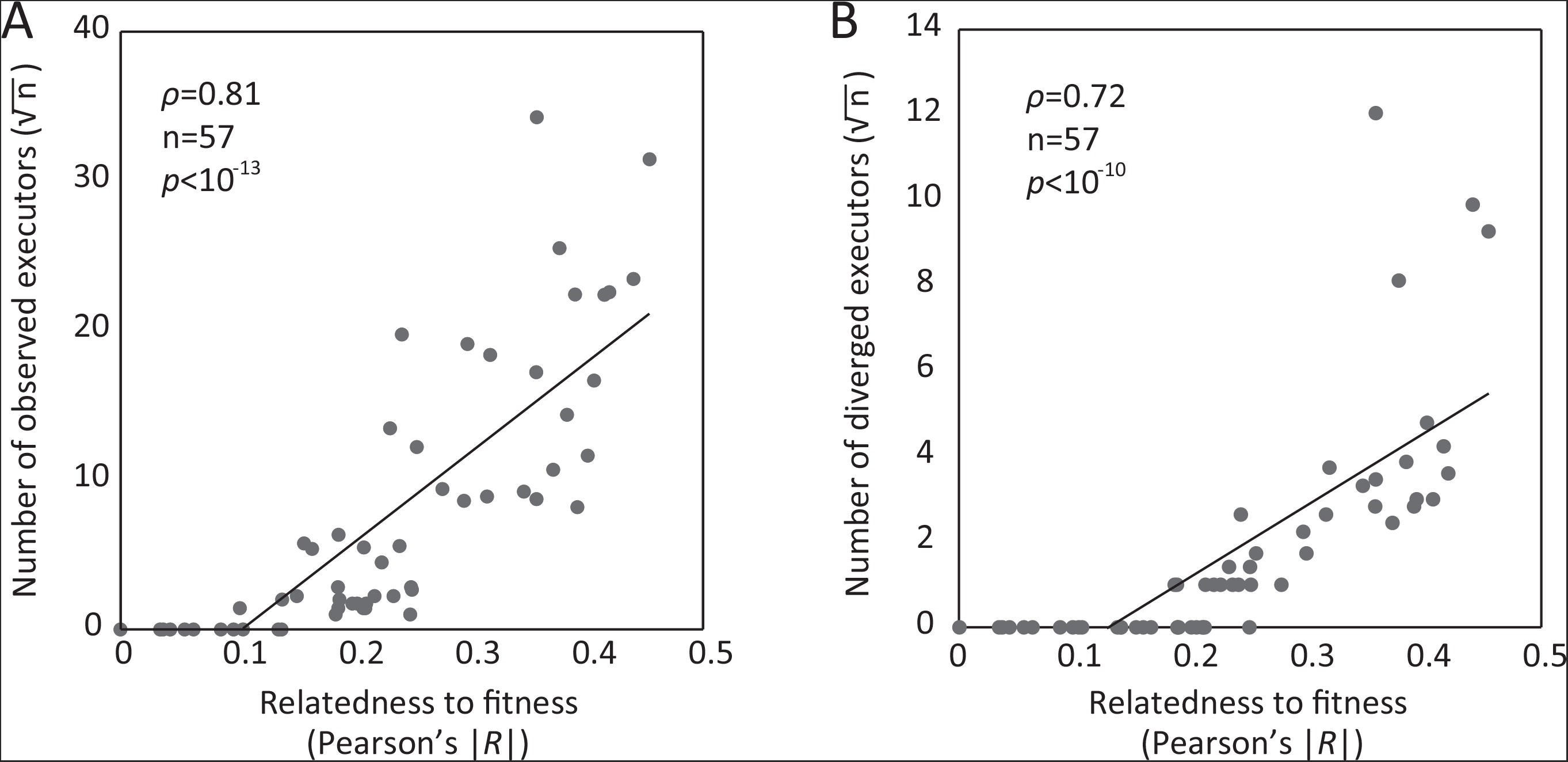
**(A)** Same as Fig. 2, except that the 57 exemplar traits are considered. **(B)** Same as Panel A, except that the observed executors present in more than one of the 57 exemplar traits are excluded.

**Fig. S4.**
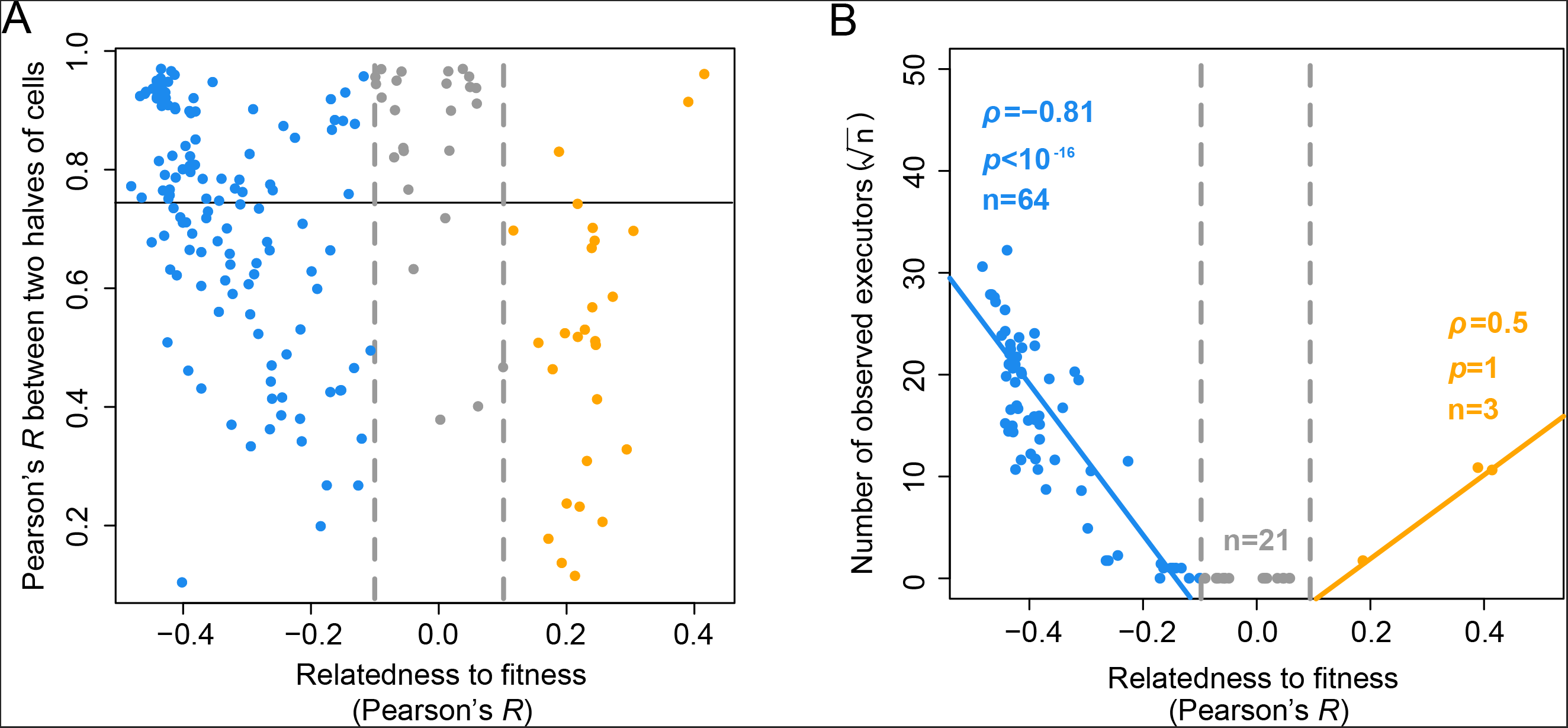
The varied quality of trait measures cannot explain the reduced number of observed executors in traits less coupled with fitness. **(A)** The y-axis shows the Pearson’s *R* of the trait values between the two halves of cells examined for each mutant, and the horizontal line marks *R* = 0.75. Each dot represents a trait, and a total of 216 traits with the information of individual cells are included. **(B)** Same as Fig. 2, except that the 88 traits with good internal consistency (*R* > 0.75) between the two halves of examined cells are considered.

**Fig. S5.**
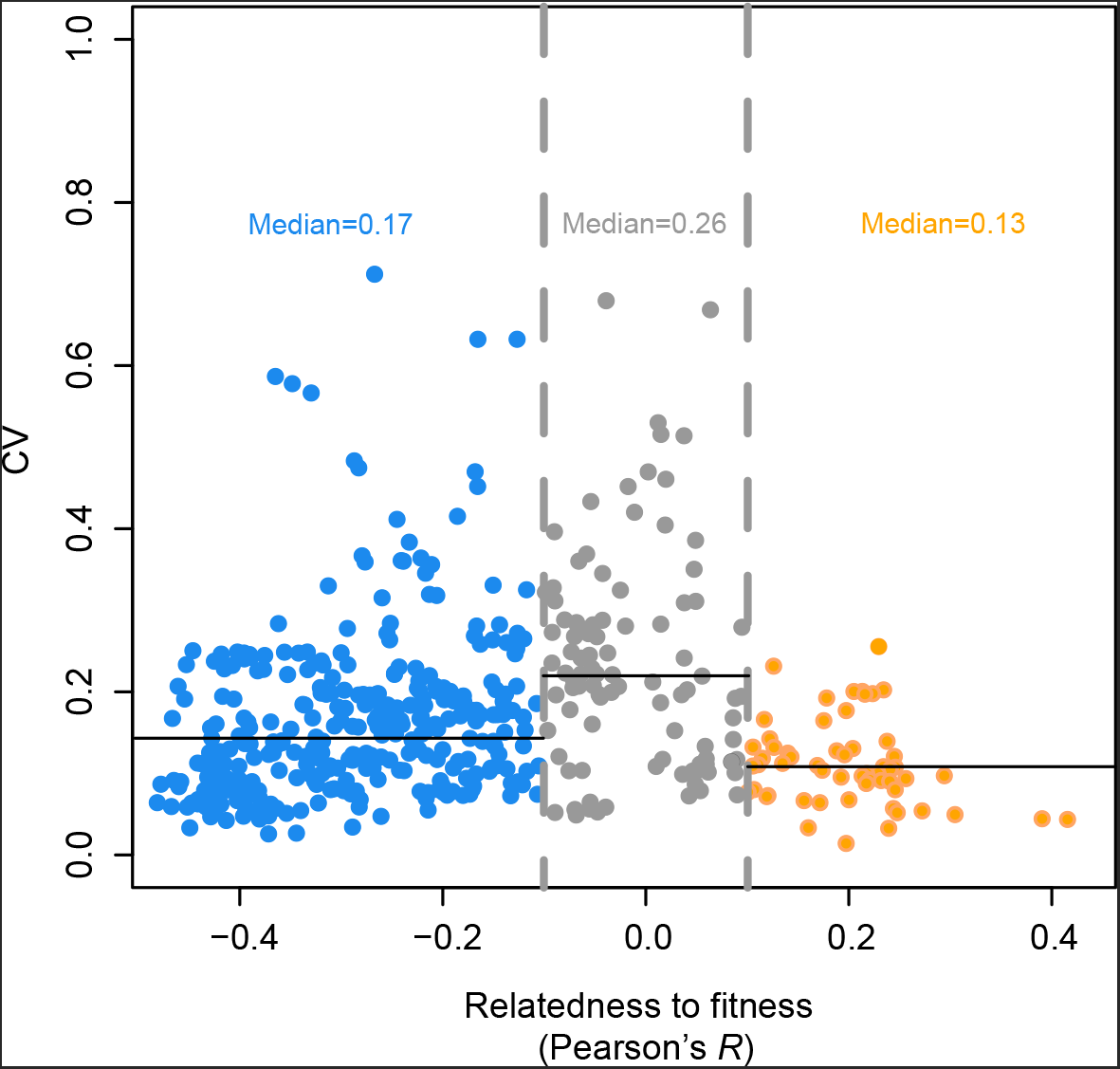
The between-mutant CV of traits with no significant correlation to fitness is not smaller than that of fitness-coupled traits, where CV stands for coefficient of variation, suggesting that the reduced number of observed executors in traits less coupled with fitness cannot be explained by the lack of variation.

**Fig. S6.**
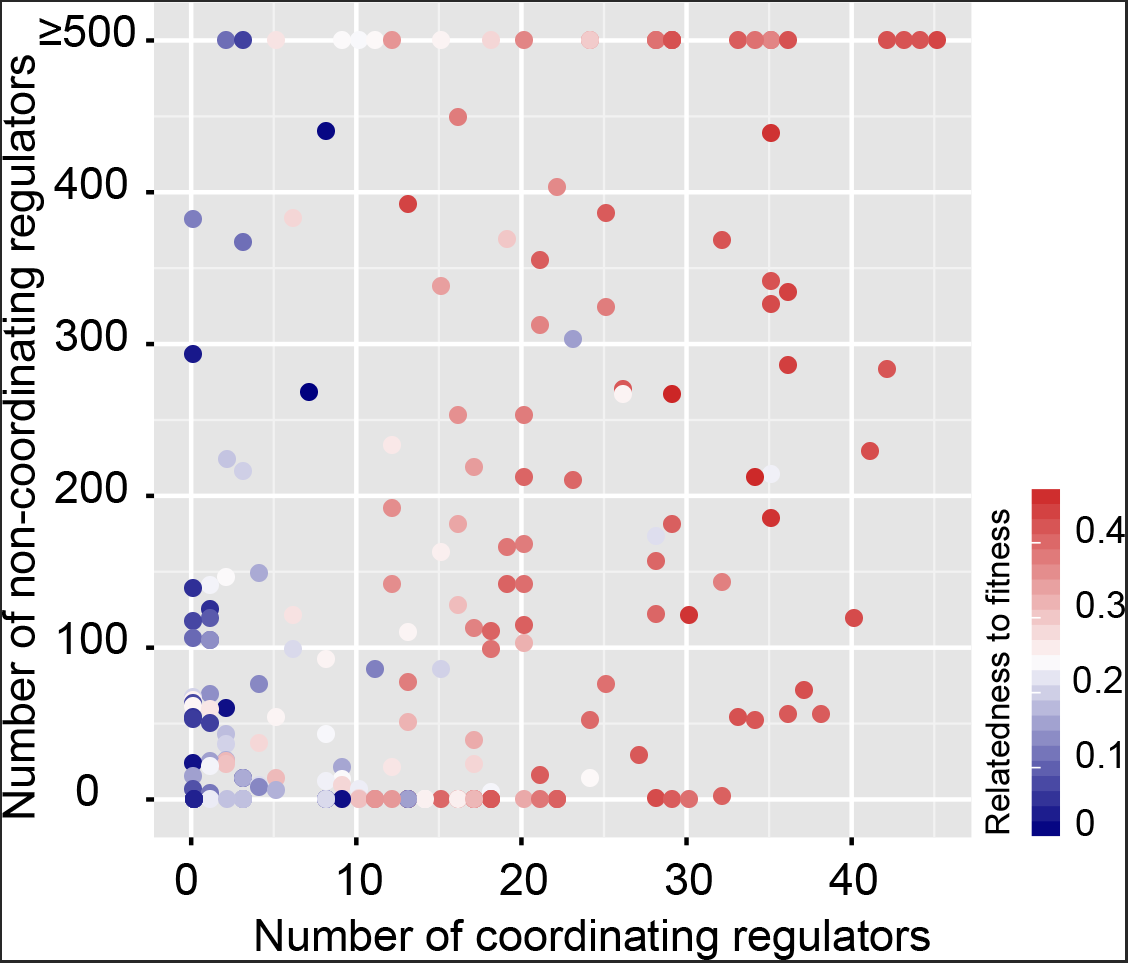
The number of coordinating regulators (outliers) and non-coordinating regulators (non-outliers) as a function of the trait relatedness to fitness. Each dot represents a trait, and a total of 216 traits are included.

**Fig. S7.**
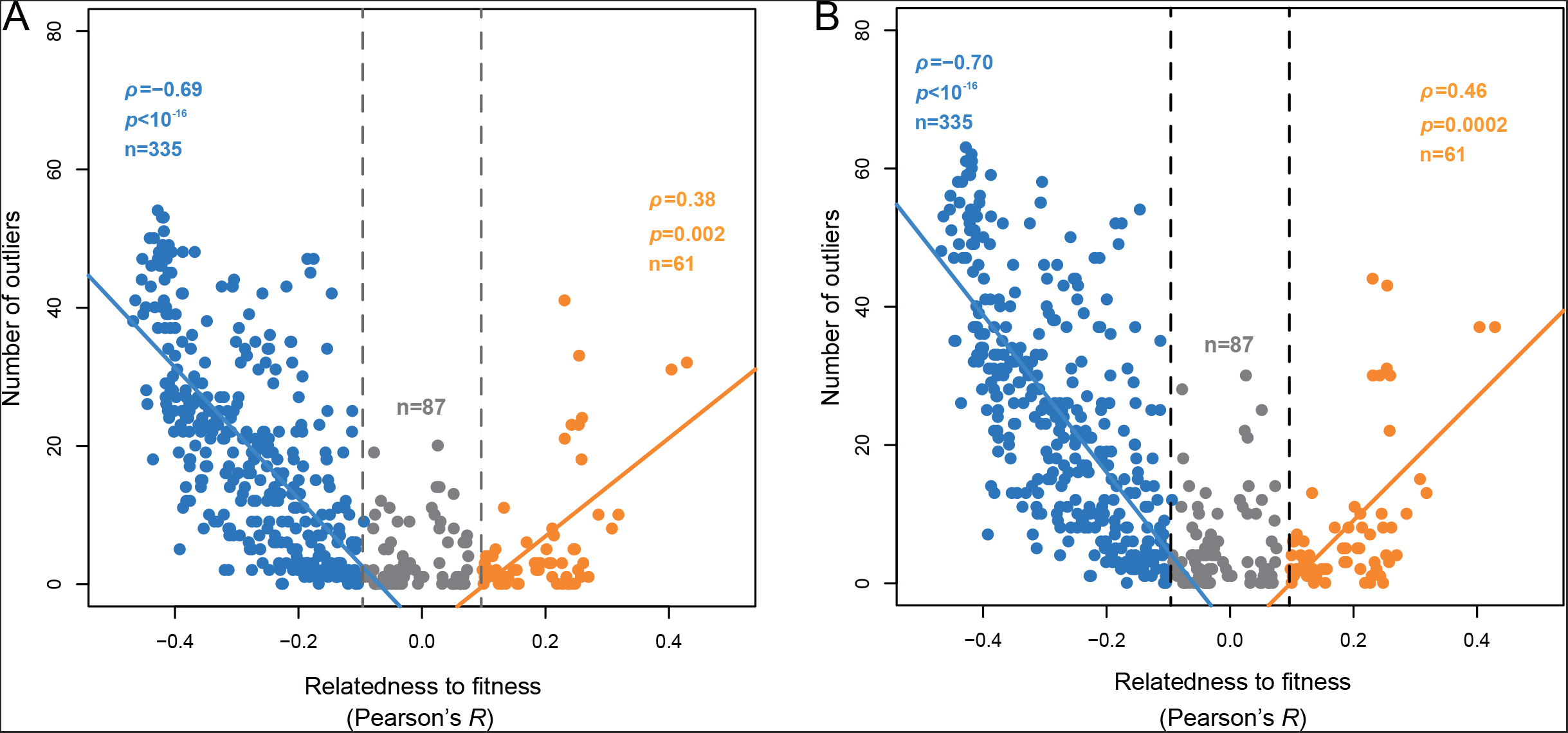
Same as Fig. 3F, except that the Z-score cutoff for identifying outliers is reduced to 4.60 **(A)** and 4.26 **(B**), corresponding to *q* smaller than 0.005 and 0.01, respectively.

**Fig. S8.**
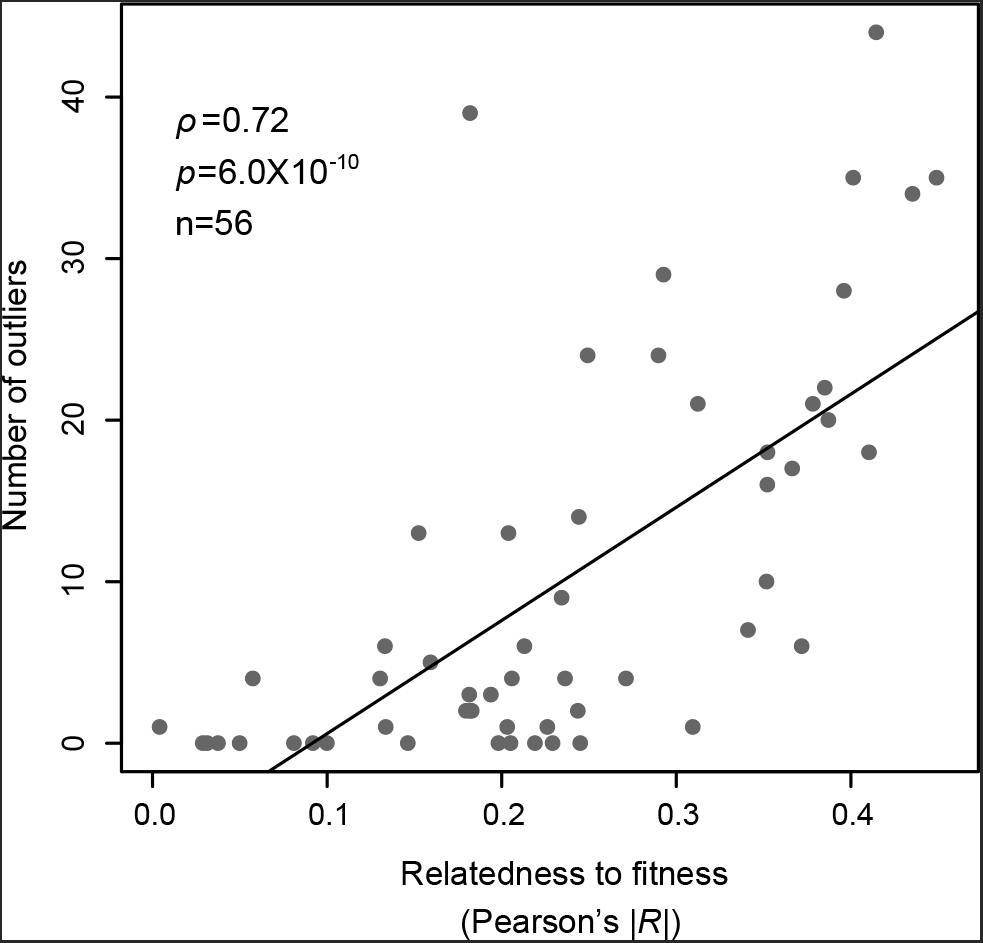
Same as Fig. 3F, except that only the 56 exemplar traits are considered, and that the absolute trait relatedness to fitness is shown at the x-axis.

**Fig. S9.**
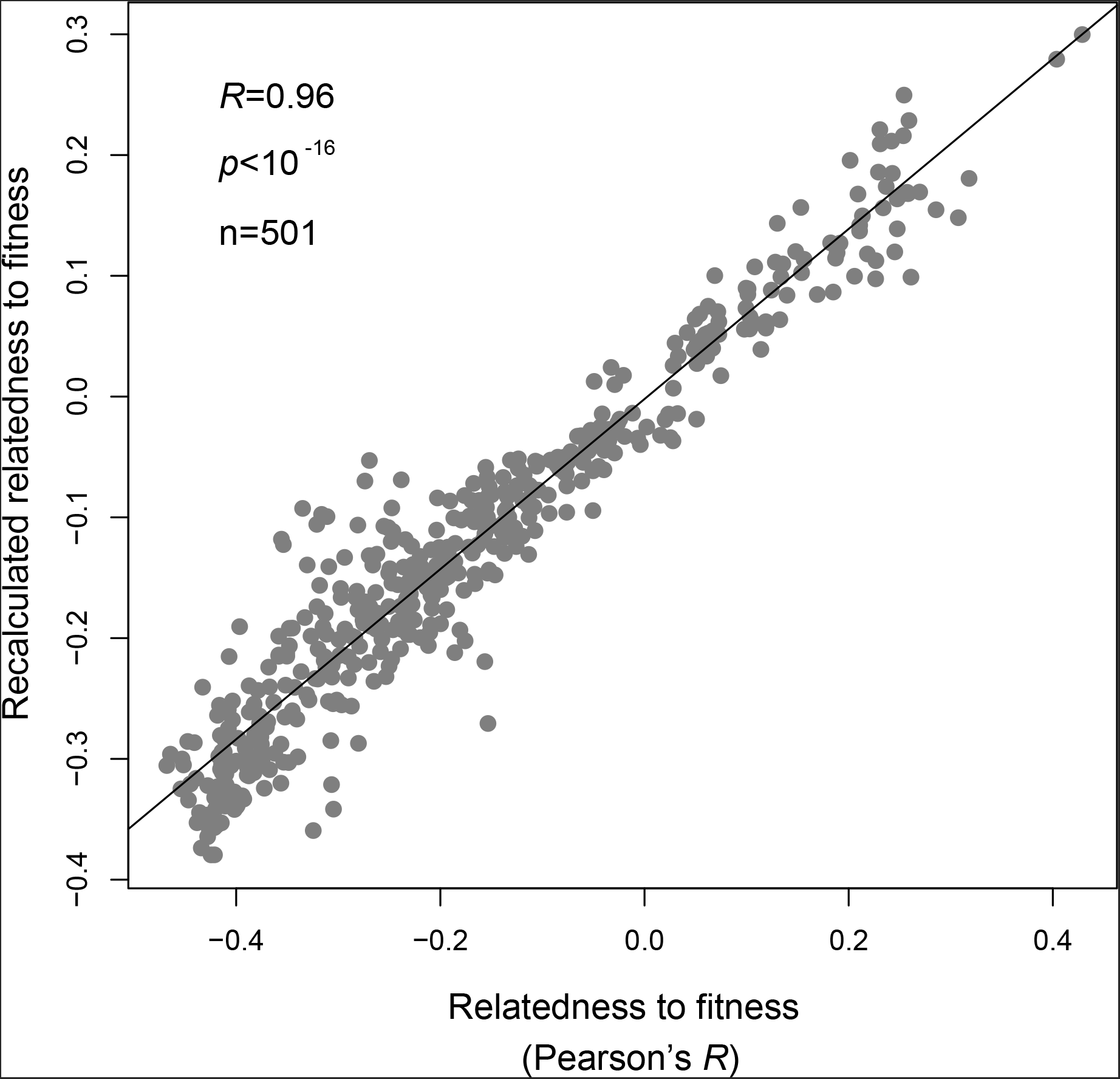
The recalculated trait relatedness to fitness (y-axis) after removing the effects of outliers is highly correlated to the original one (x-axis), indicating that fitness coupling causes the presence of outliers.

**Fig. S10.**
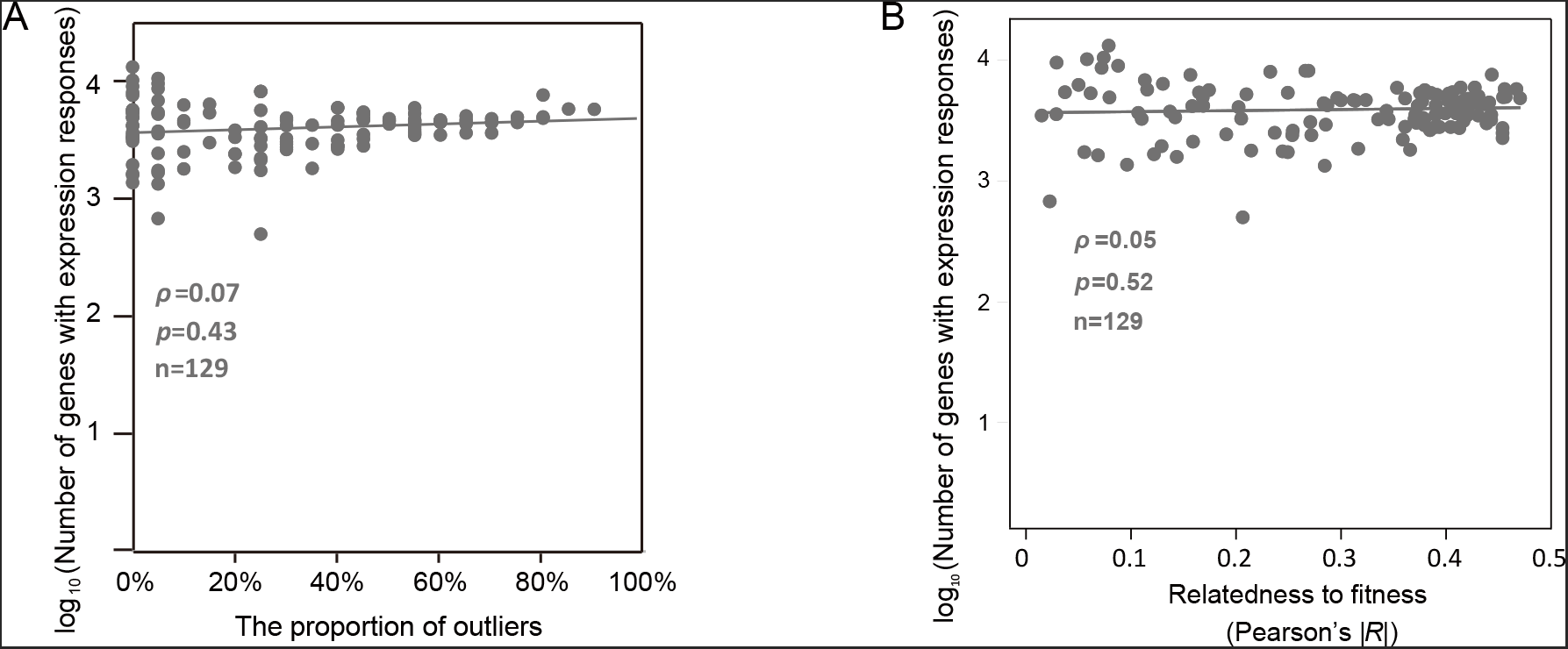
The number of genes with expression responses in mutant of any one of the selected regulators of a trait is not correlated to the proportion of outliers among the selected regulators of the trait **(A)** or the trait relatedness to fitness **(B)**. Each dot represents a trait.

**Fig. S11.**
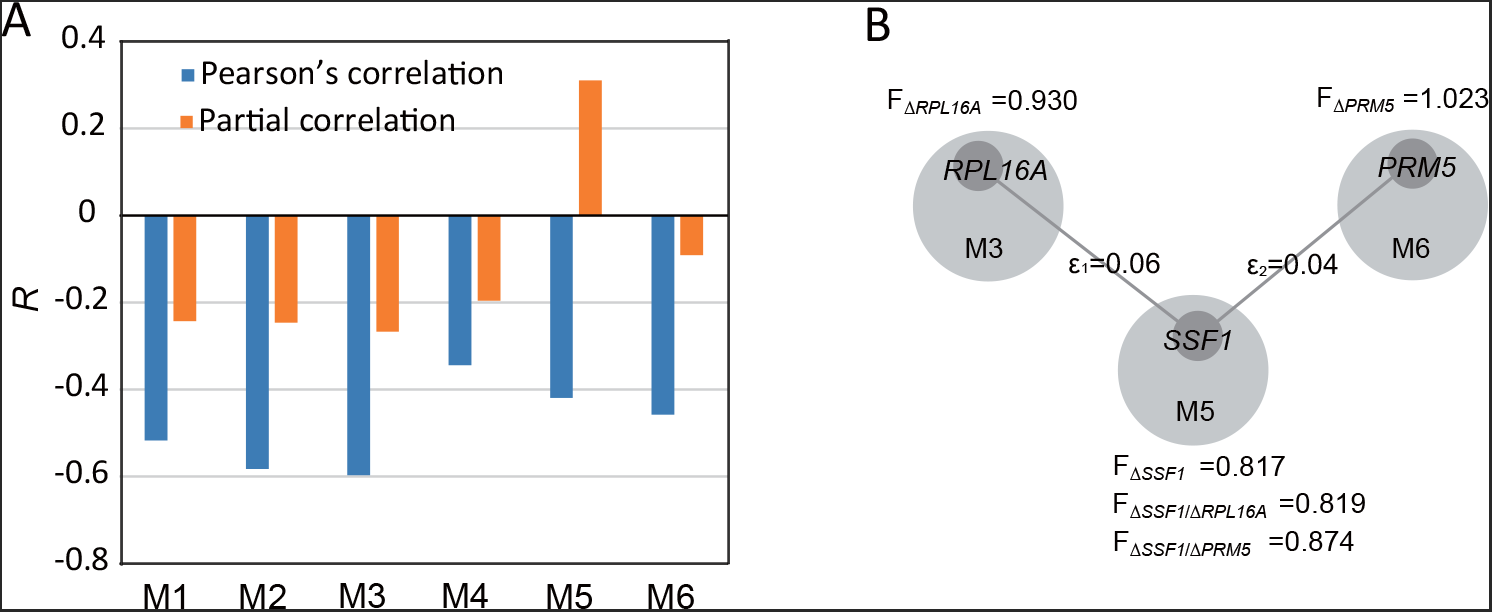
Novel mechanistic insights on yeast cell growth provided by the six executor modules. **(A)** The Pearson’s *R* between module activity and cell growth rate for each of the six executor modules, in comparison to that of the partial correlation that controls for the other five modules. **(B)** The rescuing epistasis between *SSF1* of M5 and *PRL16A* of M3 or *PRM5* of M6. F represents the relative growth rate (or fitness) of a mutant, with ε1 = F_Δ*SSF1*/Δ*PRL16A*_ – F_Δ*SSF1*_ x F_Δ*PRL16A*_ and ε2 = F_Δ*SSF1*/Δ*PRM5*_ – F_Δ*SSF1*_ x F_Δ*PRM5*_. The six executor modules are each responsible for a critical biogenesis process, including “maturation of SSU-rRNA” in module #1 (M1), “amino acid metabolism” in module #2 (M2), “translation” in module #3 (M3), “cellular respiration” in module #4 (M4), “ribosomal large subunit biogenesis” in module #5 (M5), and “cell wall organization” in module #6 (M6). Data in Fig. 5A suggest that the six modules are independent executor modules that can causally affect the yeast cell growth. This helps clarify a previous confusion with respect to the contribution of ribosome-related genes (M5) and amino acid biosynthesis genes (M2) to the yeast cell growth, which was found different between lab strains and wild strains^1^. The Pearson’s *R* between ED*M5* and the growth rate changed from -0.4 to 0.3 after controlling for the influences of the other modules. M5 represents ribosomal biogenesis, a process that consumes up to 80% of the total cell energy^2^, and its expression divergence (ED) is primarily due to the reduced gene expressions compared to the wild-type. Thus, it is likely that suppression of M5 *per se* saves energy, which promotes cell growth provided alterations of the other modules have already reduced the growth rate beneath a critical level. Consistent with these findings, deletion of *SSF1*, a member gene of M5, can be rescued by further deletion of *RPL16A*, a member gene of M3, or *PRM5*, a member gene of M6^3^. This result challenges the common belief that down-regulation of ribosomal genes reduces the cell growth rate^4-6^.

**Fig. S12.**
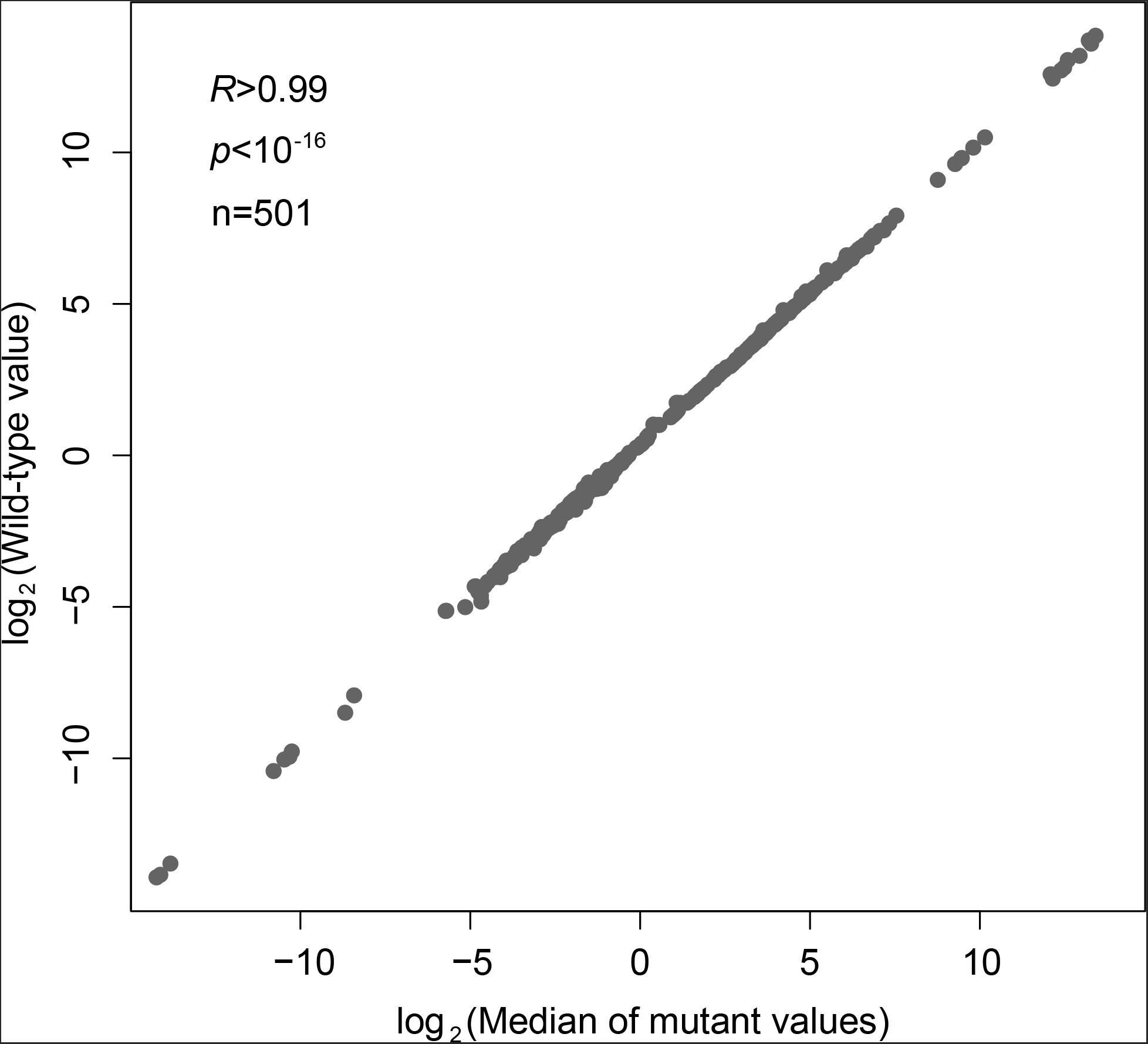
The distribution of trait values of the 4,718 mutants is bell-shaped, with the median nearly equivalent to the trait value of the wild-type for nearly all of the 501 morphological traits.

---

**Table S1.** Characterization of the six executor modules that affect the yeast cell growth.

| Module | Annotated function (GO) | *p*-value | FDR | % of genes with the GO term | Fold enrichment | Genes with the GO |
| --- | --- | --- | --- | --- | --- | --- |
| M1 | Maturation of SSU-rRNA from tricistronic rRNA transcript (SSU-rRNA, 5.8S rRNA, LSU-rRNA) | 2.76X10<sup>-32</sup> | 1.53X10<sup>-28</sup> | 20/31=64.5% | 46.07 | YCR057C YOR056C YGR090W YDR324C YDR449C<br>YJL191W YJL010C YBR247C YMR229C YLR222C<br>YJR002W YKL078W YPL081W YIL019W YDL014W<br>YOR004W YDR398W YPL012W YER082C YDL153C |
| M2 | Cellular amino acid and derivative metabolic process | 1.32X10<sup>-06</sup> | 6.49X10<sup>-05</sup> | 9/24=37.5% | 7.50 | YGR061C YDR502C YHR019C YGR155W YBR121C<br>YKL104C YBL076C YGL026C YLR180W |
| M3 | Translation | 3.04X10<sup>-06</sup> | 9.31X10<sup>-05</sup> | 12/19=63.1% | 4.10 | YNL301C YHL033C YNL096C YBR031W YPL090C<br>YPR041W YNL178W YLR287C-A YDR012W<br>YMR194W YLL045C YIL133C |
| M4 | Cellular respiration | 4.82X10<sup>-17</sup> | 1.45X10<sup>-15</sup> | 16/45=35.5% | 18.68 | YKL141W YGR183C YDL067C Q0250 YLL041C<br>YKR046C YJL166W YMR256C YOR065W YPR191W<br>YGL191W YBL045C YHR051W YDR178W YEL024W<br>YDR529C |
| M5 | Ribosomal large subunit biogenesis | 5.07X10<sup>-21</sup> | 3.28X10<sup>-19</sup> | 14/26=53.8% | 44.83 | YDR060W YOR294W YMR290C YLL008W<br>YPL093W YHR197W YBR142W YNL182C YHR052W<br>YGR103W YHR066W YHR085W YGR245C YCR072C |
| M6 | Cell wall organization | 2.40X10<sup>-06</sup> | 7.12X10<sup>-05</sup> | 23/190=12.1% | 3.03 | YKR100C YOL030W YNL322C YNL047C YGR279C<br>YDR055W YNL066W YNL283C YKL163W YBR180W<br>YNL192W YKL129C YBR023C YJL159W YOR247W<br>YLR390W-A YMR200W YGR189C YMR215W<br>YLR380W YMR104C YIL117C YDR077W |

